# Genetic factors influencing a neurobiological substrate for psychiatric disorders

**DOI:** 10.1101/774463

**Authors:** Till F. M. Andlauer, Thomas W. Mühleisen, Felix Hoffstaedter, Alexander Teumer, Katharina Wittfeld, Anja Teuber, Céline S. Reinbold, Dominik Grotegerd, Robin Bülow, Svenja Caspers, Udo Dannlowski, Stefan Herms, Per Hoffmann, Tilo Kircher, Heike Minnerup, Susanne Moebus, Igor Nenadić, Henning Teismann, Uwe Völker, International FTD-Genomics Consortium (IFGC), The 23andMe Research Team, Amit Etkin, Klaus Berger, Hans J. Grabe, Markus M. Nöthen, Katrin Amunts, Simon B. Eickhoff, Philipp G. Sämann, Bertram Müller-Myhsok, Sven Cichon

## Abstract

A retrospective meta-analysis of magnetic resonance imaging voxel-based morphometry studies proposed that reduced gray matter volumes in the dorsal anterior cingulate and the left and right anterior insular cortex – areas that constitute hub nodes of the salience network – represent a common substrate for major psychiatric disorders. Here, we investigated the hypothesis that the common substrate serves as an intermediate phenotype to detect genetic risk variants relevant for psychiatric disease. To this end, after a data reduction step, we conducted genome-wide association studies of a combined common substrate measure in four population-based cohorts (n=2,271), followed by meta-analysis and replication in a fifth cohort (n=865). After correction for covariates, the heritability of the common substrate was estimated at 0.50 (standard error 0.18). The top single-nucleotide polymorphism (SNP) rs17076061 was associated with the common substrate at genome-wide significance and replicated, explaining 1.2% of the common substrate variance. This SNP mapped to a locus on chromosome 5q35.2 harboring genes involved in neuronal development and regeneration. In follow-up analyses, rs17076061 was not robustly associated with psychiatric disease, and no overlap was found between the broader genetic architecture of the common substrate and genetic risk for major depressive disorder, bipolar disorder, or schizophrenia. In conclusion, our study identified that common genetic variation indeed influences the common substrate, but that these variants do not directly translate to increased disease risk. Future studies should investigate gene-by-environment interactions and employ functional imaging to understand how salience network structure translates to psychiatric disorder risk.

## Introduction

Numerous studies have identified regional differences in the brain structure of psychiatric patients and described both transdiagnostic and disorder-specific processes of gray matter (GM) reduction in patients (1–8). One of these reports was the large retrospective meta-analysis of 193 studies by Goodkind *et al*. that compared 7,381 psychiatric patients from six diagnostic groups (schizophrenia, bipolar disorder (BD), major depressive disorder (MDD), addiction, obsessive-compulsive disorder, and anxiety) with 8,511 psychiatrically healthy controls using voxel-based morphometry (VBM) from structural magnetic resonance imaging (MRI) (1). Across all diagnoses, they found that GM volumes were lower in the left and right anterior insular cortices (AIC) and the dorsal anterior cingulate cortex (dACC). Subsequentially, they performed structural and functional connectivity analyses and confirmed that these three regions were tightly connected and represent hub nodes of the salience network (1, 9, 10): This network serves stimulus selection, controls the focus of attention, is involved in the selection of goal-directed behavior and in saliency detection of exogenous or internal cues (9,10,11). Independent studies indicate that functional differences in salience processing in these brain regions are associated with several neuropsychiatric disorders and their progression (11). Eventually, Goodkind and colleagues hypothesized that lower GM of this network represents a common neurobiological substrate for psychiatric disorders (1).

However, the etiology of the common substrate reductions has not been investigated so far and remains unclear. One possible explanation involves the loss of GM at disease manifestation and during the further course of disease, implying a regionally specific vulnerability towards a degenerative process – similar to known neurodegenerative disease entities (12, 13). An alternative explanation implies that reduced GM exists before disease onset, shaped by genetic or early environmental influences such as childhood adversity (14): Here, premorbid structural abnormalities of the salience network could increase a subject’s vulnerability to psychiatric disease. More recently, structural salience network integrity was reported to mediate between polygenic risk for schizophrenia and auditory hallucinations (15). A third explanation involves brain-aging processes that occur in a network-dependent way and often with a strong non-linear component (16–18): Here, accelerated aging could increase the disease risk over the lifespan by genetic or environmental factors. All three explanation models might apply in parallel and lead to combined effects at the morphological level.

Many studies have analyzed genetic risk factors for psychiatric disorders such as schizophrenia, BD, and MDD (19). These disorders show substantial heritability (20) and are genetically correlated with each other (21, 22). Genome-wide association studies (GWAS) identified singlenucleotide polymorphisms (SNPs) contributing risk for several psychiatric disorders, suggesting pleiotropy and partially overlapping etiologies (22, 23). Imaging genomics is a growing discipline that exploits imaging-based measures to explore the genetic basis of brain organization (24). The clinical value of this concept to detect risk variants for psychiatric disease, however, depends on a detectable correlation between the intermediate phenotype and the clinical level. Following this line of thought, the CS suggested by Goodkind and colleagues is a promising intermediate phenotype, particularly due to its transdiagnostic effects.

The present study aimed to identify genetic variants influencing the substrate in the general population. As a conceptual decision, patient cohorts were not included in our genetic analyses to avoid any interference with secondary disease effects on the common substrate, such as treatment effects or other disease-related epiphenomena. Our imaging analyses involved a prospective, harmonized VBM preprocessing protocol applied to high-resolution structural MRI data of five population-based cohorts. To account for the network character of the three common substrate regions, we combined them into a single marker using principal component analysis. We analyzed the first principal *component of the common substrate* (CCS) of the population-based cohorts through GWAS, followed by meta-analysis. As our main result, we identified a novel genetic locus significantly associated with the CCS. In a series of secondary analyses, we characterized the genetic relationship between the CCS and risk for psychiatric disorders and investigated a potentially modulating role of age.

## Methods and Materials

### Sample description

For the GWAS, 3,136 individuals from five population-based cohorts were pooled. Four cohorts were used in the discovery (1000BRAINS (25), n=539; CONNECT100 (26), n=93; BiDirect (27), n=589; SHIP-2 (28), n=1,050; total n=2,271) and the second-largest cohort available was used in the replication stage (SHIP-Trend (28), n=865). For follow-up analyses, three psychiatric patient/control cohorts with 1,978 patients and 1,375 controls were used, BiDirect (n=582 MDD patients; n=311 healthy controls (29)), MPIP (n=385 MDD patients; n=197 healthy controls (30, 31)), and FOR2107 (n=769 MDD, n=127 BD, n=72 schizophrenia, and n=43 schizoaffective patients; n=867 healthy controls (32, 33)). The BiDirect cohort is a prospective observational study (27). The Max Planck Institute of Psychiatry (MPIP) cohort represents subsamples of the Munich Antidepressant Response Signature study (MARS), an observational study on psychiatric in-patients treated for MDD (30), and the recurrent unipolar depression study, a cross-sectional case/control imaging genetics study (31) (see (5,6) for diagnostic instruments). Probands were recruited in the area of Münster and underwent a structured clinical interview for DSM-IV axis I disorders and all MDD patients received treatment for acute depression (27). FOR2107 is an ongoing multi-center study recruiting from the areas of Marburg and Münster in Germany (32). All subjects underwent a structured clinical interview for DSM-IV axis I disorders (SCID-I), administered by trained clinical raters. Basic demographic characteristics of the cohorts can be found in Supplementary Tables S1 and S2. The studies were approved by the local ethics committees; all participants provided written informed consent.

### VBM preprocessing and extraction of regional and total GM volumes

VBM-like preprocessing with MATLAB-based SPM (version 8, https://www.fil.ion.ucl.ac.uk/spm/software/spm8/) and the VBM8 toolbox (version r445, http://dbm.neuro.uni-jena.de/vbm8/) were used to process all T1-weighted images (n=3,136 for the GWAS and n=3,361 for the patient/control analyses). Processing was performed locally by the participating sites and comprised the following steps: (i) spatial registration to a reference brain in MNI152 space, (ii) segmentation of T1-weighted images into GM, white matter, and cerebrospinal fluid by a three-step algorithm implemented in the vbm8 toolbox, (iii) bias correction of intensity non-uniformities, (iv) application of the diffeomorphic normalization algorithm DARTEL for iterative linear and non-linear spatial normalization of the GM and white matter maps to MNI space (IXI555 template) (34), (v) non-linear-only Jacobian modulation to correct for linear global scaling effects while preserving local GM volumes. The quality of processed GM segments in MNI space was assessed using the “Check sample homogeneity using covariance” function in VBM8. Three spatially disjunct regional GM volumes, based on binarized versions of the joint result areas from the study by Goodkind *et al*. (1), and total GM volume were extracted.

Extracted GM volumes were, separately for each cohort, corrected for age, age^2^, and sex in multiple linear regression models. Handedness was used as an additional covariate for 1000BRAINS, CONNECT100, and BiDirect, coil type for the MPIP sample, as well as body coil type and site for FOR2107. Residuals of these regional volume regression models were combined using principal component analysis (PCA) to create a single measure, which we named the CCS (Fig. 1).

**Figure 1:**
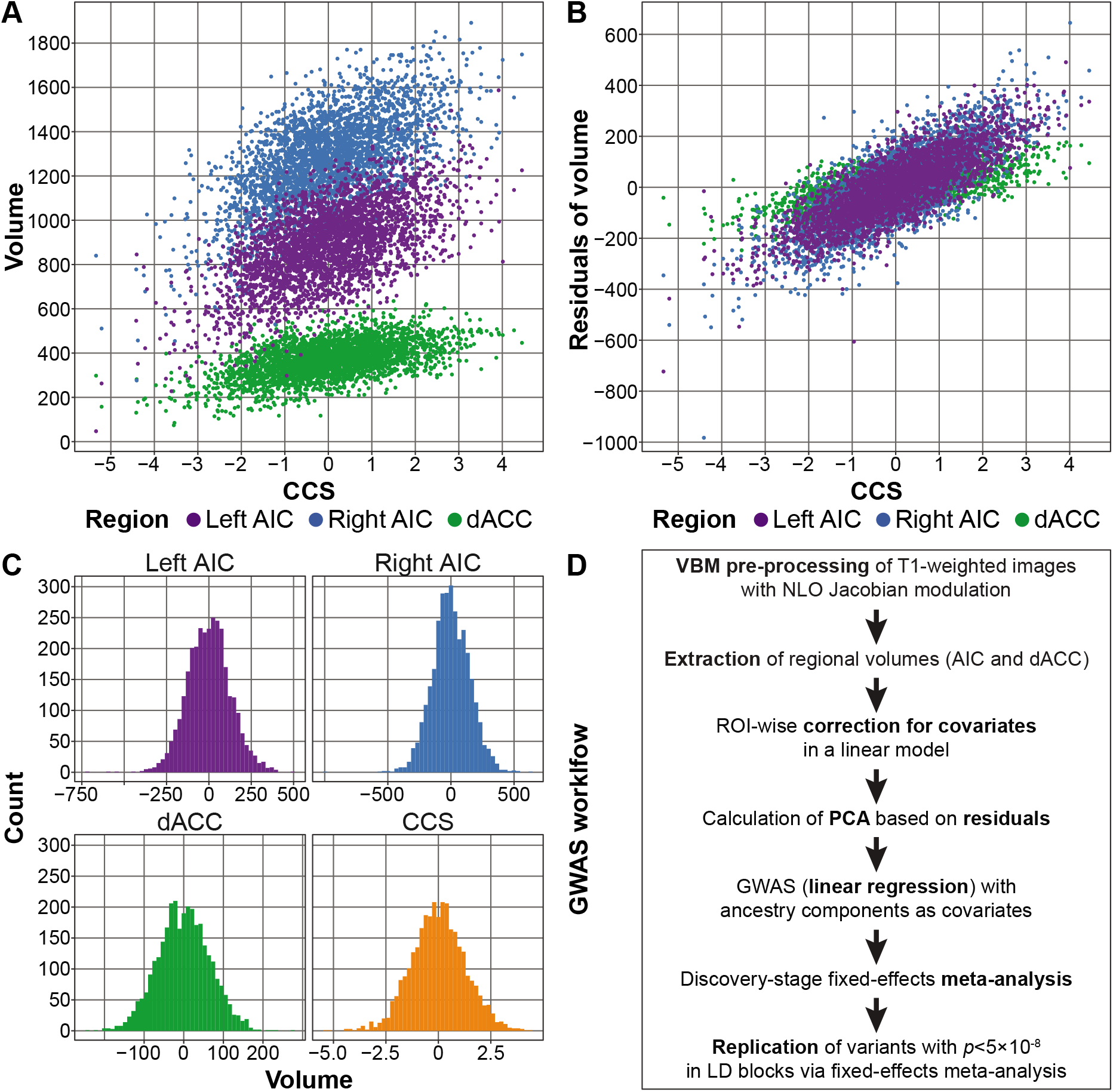
Generation of the component of the common neurobiological substrate (CCS) and genome-wide association study (GWAS) analysis workflow. **A-B:** Comparison between the CCS and the three individual volumes (A) and the residuals of the three volumes after correction for covariates (B). AIC = anterior insula cortex; dACC = dorsal anterior cingulate cortex **C:** Histograms of the three extracted volumes and the CCS. Note that A-C show combined data from all five GWAS cohorts. Correlations were left and right AIC: *r*=0.65, left AIC and dACC: *r*=0.52, right AIC and dACC: *r*=0.46. **D:** GWAS analysis workflow. All measures were extracted using NLO-based Jacobian modulation. All GM volumes were corrected for age, age^2^, and sex as covariates; handedness was used as an additional covariate for the three samples 1000BRAINS, CONNECT100, and BiDirect. **PCA =** principal component analysis; **LD =** linkage disequilibrium.

### Genotyping, quality control, and imputation

DNA extraction and genome-wide genotyping were conducted as described before (28, 31, 35–37). Genotyping was carried out on different Illumina and Affymetrix microarrays (see the Supplementary Methods and Supplementary Table S3). Quality control (QC) and imputation were conducted separately for each genotyping batch, using the same protocols, in PLINK, *R*, and XWAS (38, 39). Genotype data were imputed to the 1000 Genomes phase 1 reference panel using SHAPEIT and IMPUTE2 (40–41), as described in the Supplementary Methods and previously (42). The population substructure of all five GWAS cohorts is shown in Supplementary Figure S1.

### Heritability estimation and GWAS

The SNP-based heritability of the CCS was estimated using genome-wide complex trait analysis (GCTA) on a combined sample of the imputed data from all five cohorts (43) (see the Supplementary Methods). GWAS was conducted separately per cohort using PLINK with ancestry components as covariates. Variants on the X chromosome were analyzed separately by sex, followed by *p* value-based meta-analysis to allow for different effect sizes per sex. A two-stage design was implemented for the GWAS, using four cohorts as the discovery sample and SHIP-Trend as an independent replication sample. The cohorts were combined with fixed-effects meta-analysis using METAL (44). There was no indication for genomic inflation of the GWAS test statistics in the single cohorts or the meta-analysis (λ_1000_=1.01, see Supplementary Table S4 and Supplementary Figure S2).

Linkage disequilibrium (LD) was analyzed using the European 1000 Genomes CEU population in *LDmatrix* (45). The two SNPs that showed the most robust genome-wide support (*p*<5×10^−8^) for an association in the discovery stage and were partially independent of each other (LD *r^2^*<0.5 with more strongly associated variants) were carried forward to the replication stage. Here, a one-sided *p*-value <α=0.05/2 (correcting for two LD-independent variants) was considered as a successful replication. See the Supplementary Methods for a detailed description of heritability and GWAS methods.

### Gene-set analyses

Gene-set analyses were conducted on the meta-analysis of the discovery- and replication-stage GWAS, using 674 RE-ACTOME gene sets containing 10–200 genes curated from *MsigDB 6.2* (46). Only SNPs within gene boundaries were mapped to RefSeq genes (0 bp window). Analyses were conducted in MAGMA v1.07 using both mean- and top-SNP gene models (47) and in MAGENTA v2 using a top-SNP approach (48). Here, false discovery rates were calculated to correct for multiple testing.

### Comparison to published GWAS of psychiatric disorders and polygenic score analyses

For genome-wide comparisons between our GWAS meta-analysis and published GWAS of psychiatric disorders, summary statistics from the following Psychiatric Genomics Consortium (PGC) GWAS were used: cross-disorder 2019 (22), BD 2019 (49), MDD 2018 (with 23andMe) (50), and schizophrenia 2014 (51). For additional comparisons, the following GWAS were used: IFGC behavioral frontotemporal dementia (bvFTD) 2014 (52), longevity 85/90 2014 (53), and three different GWAS from 2017 on epigenetic accelerated aging (EAA) (54): accelerated aging in all examined brain regions, accelerated aging in prefrontal cortex, and neuronal proportion in the prefrontal cortex.

To further characterize the relationship between the CCS and risk for psychiatric disorders, we ran four analyses using GWAS summary statistics from published PGC studies, following a published, well-acknowledged workflow (55). Polygenic scores (PGSs) were calculated and analyzed in *R* using imputed genetic data (56, 57). Here, we used the PGC GWAS as training and our population-based GWAS cohorts as test data. Furthermore, we also calculated PGS using the CCS GWAS summary statistics as training and the patient/control cohorts as test data. We ran LD score regression (LDSC) comparing the genetic correlation of published GWAS to the CCS GWAS summary statistics with standard settings (58, 59). We analyzed whether the order of SNPs ranked by their association strength was random between studies using rank-rank hypergeometric overlap (RRHO) tests (60). For this analysis, variants were LD-pruned in the 1000 Genomes phase 3 EUR subset (61). Binomial sign tests were conducted on LD-clumped variants in *R* (*binom.test*) to analyze whether SNPs associated with the CCS at either *p*<0.05 or *p*<1×10^−5^ showed the opposite direction of effects in other GWAS more often than expected by chance. For additional details on these analyses, see the Supplementary Methods.

### Secondary analyses of age interaction effects

We explored the possibility that the original VBM studies entering the meta-analysis of Goodkind *et al*. (1) picked up age-by-diagnosis effects by analyzing patient/control cohorts and by verifying that our main genetic association was not age-dependent. We performed secondary analyses that probed (a) the possibility of ‘accelerated aging’ of the CCS phenotype in psychiatric disorders and (b) the possibility of heterogeneity of the SNP effect across different age ranges. To allow for a valid interpretation of the age-interaction terms, we conducted a Gram-Schmidt orthonormalization of age (first term) and age^2^ (second term) in *R* v.3.5.2, using the function *QR* of the package *mat-lib*.

## Results

### Combination of the three brain regions

To analyze a combined measure of the published common neurobiological substrate for psychiatric disorders (1), we combined the volumes of the left AIC, right AIC, and dACC by PCA. The first principal component, which we refer to as the CCS, explained 66.5% of the phenotypic variance of the three volumes and 55.4% after correction of the volumes for covariates (Fig. 1, Supplementary Methods).

### Heritability of the CCS

After correction for covariates, the CCS showed a SNP heritability estimate of *h*^2^_*g*_=0.50 (standard error (SE)=0.18; *p*-value=0.0033).

### GWAS of the CCS

In the discovery-stage GWAS (Supplementary Fig. S2A and Supplementary Table S4), twelve SNPs on chromosome *5q35.2* showed genome-wide significant associations with the CCS (significance threshold *p*<5×10^−8^; Fig. 2A and Supplementary Table S5). Most of these variants were highly correlated with each other (Supplementary Table S6). The two partially LD-independent SNPs (pairwise LD *r^2^*=0.267 in CEU samples) with the most robust support for an association were analyzed further (Fig. 2B). Of these, the minor allele T of the SNP rs17076061 (frequency in our GWAS cohorts: 0.36, Fig. 2C) was significantly associated in the replication cohort in the same direction (discovery: β=-0.22 standard deviations (SE=0.04), *p*=1.51×10^−8^; replication: β=-0.15 (SE=0.07), one-sided *p*=9.91×10^−3^) and was also the top-associated variant in the genome-wide meta-analysis of discovery and replication samples (β=-0.21 (SE=0.03), *p*=1.46×10^−9^; Fig. 2D, Supplementary Table S5, Supplementary Figs. S2B, S3 and S4). SNP rs17076061-T was associated with the CCS at genome-wide significance but not with the three single region volumes or the whole-brain GM volume (Table 1). After z-score transformation to allow effect size comparisons, the effect size was larger for the CCS (−0.159 standard deviations (SD) than for the total GM (−0.099 SD).

**Table 1:** Association results from the genome-wide meta-analysis of discovery and replication samples in different gray matter (GM) regions. For this comparison, all measures were centered and scaled using z-score transformation before the analysis to make the effect sizes of the different measures comparable. The unit of the effect sizes are thus standard deviations (SD). Accordingly, the CCS coefficients shown here differ from the ones presented in Fig. 2 and Supplementary Table S5. The effect size refers to the minor allele T. All measures were extracted using non-linear only (NLO)-based Jacobian modulation. **AIC** = anterior insula cortex; **dACC** = dorsal anterior cingulate cortex; **SE** = standard error.

| **rs17076061** | **Effect size** | **SE** | **$p$-value** |
| --- | --- | --- | --- |
| CCS | -0.159 | 0.026 | $1.41\times10^{-9}$ |
| Left AIC | -0.142 | 0.026 | $7.00\times10^{-8}$ |
| Right AIC | -0.124 | 0.026 | $2.63\times10^{-6}$ |
| dACC | -0.083 | 0.026 | $1.77\times10^{-3}$ |
| Total gray matter | -0.099 | 0.026 | $1.85\times10^{-4}$ |

**Figure 2:**
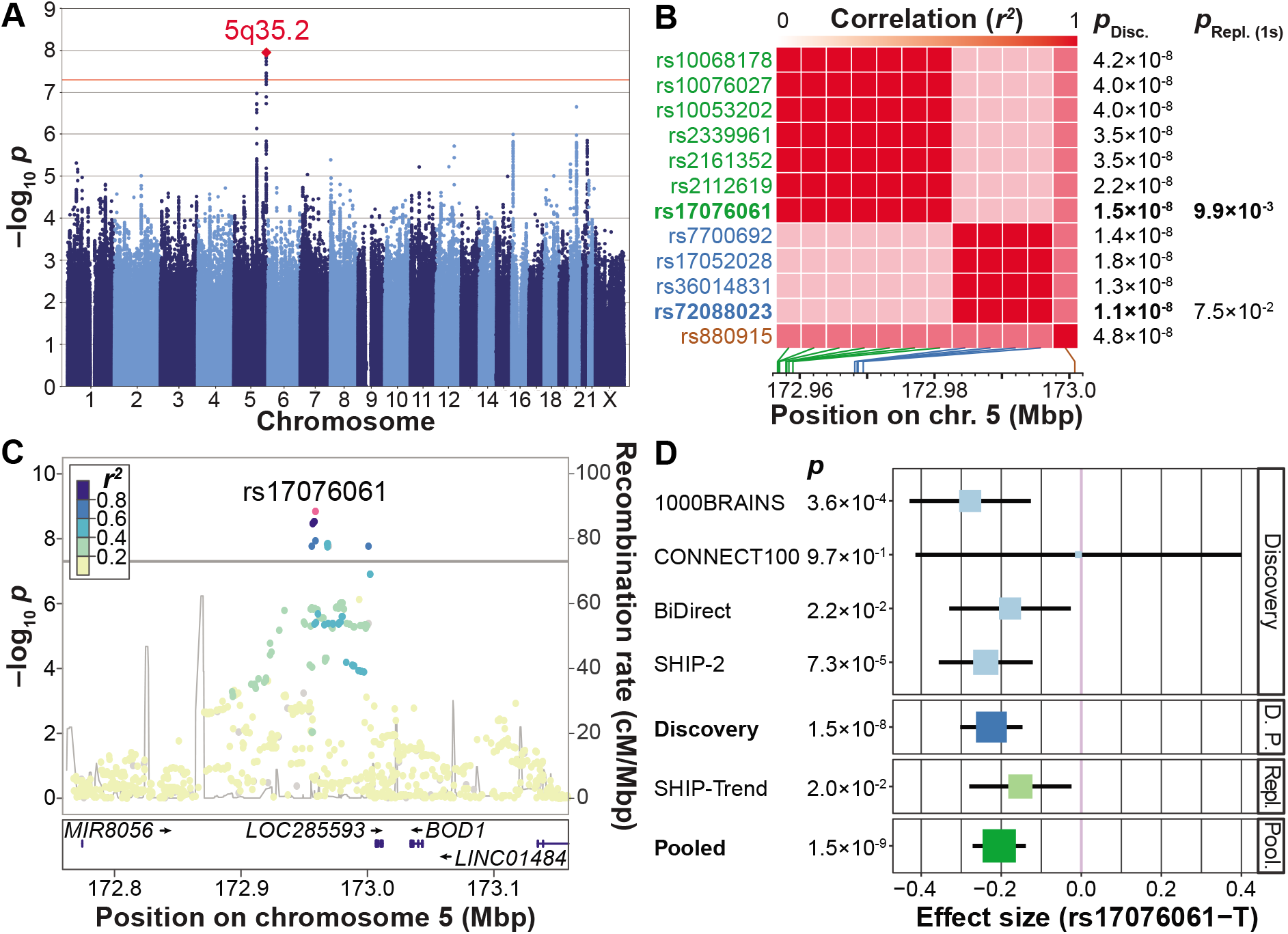
Presentation of the genome-wide association study (GWAS) results. **A:** Manhattan plot showing the strength of evidence for an association (*p*-value) in the discovery stage component of the common neurobiological substrate (CCS) GWAS. Each variant is shown as a dot, with alternating shades of blue according to chromosome; the top-associated locus *5q32.2* is labeled with a red diamond. The red line marks the genome-wide significance level. **B:** Matrix of the pairwise linkage disequilibrium (LD) pattern (1000 Genomes population CEU) between the twelve variants that reached genome-wide significance in the discovery GWAS. The two variants rs17076061 and rs72088023 (*r^2^*=0.267) showed the strongest support for an association in their respective LD blocks and were analyzed in the replication stage. All other variants had pairwise LD>0.5 with either of these two variants, their association strengths are provided for comparison only. *P*_bisc_.: discovery stage GWAS *p*-value; *p*_Repi.(1s)_: one-sided *p*-value in the replication cohort; Mbp: mega base pair. **C:** Regional association plot of the top-associated locus after pooled analysis of the discovery stage GWAS and the replication sample. The dot color indicates LD with the lead variant (rs17076061; pink). Gray dots represent signals with missing LD *r^2^* values. cM: centimorgan. **D:** Forest plot of the pooled analysis of the replicated variant rs17076061 in discovery and replication cohorts. D. P.: pooled analysis of discovery stage cohorts; Repl.: replication; Pool.: pooled analysis of the discovery GWAS and the replication cohort SHIP-Trend.

### Gene-set analyses

In two separate gene-set analyses using GWAS meta-analysis results, four pathways were significantly associated with the CCS. The top-associated pathway in both analyses (MAGMA: adjusted *p*=2.2×10^−3^; MAGENTA: false discovery rate *q*=2.4×10^−3^) was “SEMA3A-Plexin repulsion signaling by inhibiting Integrin adhesion” (https://www.reactome.org/content/detail/R-HSA-399955). Please see Supplementary Tables S7 and S8 for the full results of these analyses.

### Comparison of the top GWAS SNP and the genetic architecture of the CCS with genetic risk for disease

To investigate whether rs17076061 is associated with risk for common psychiatric disorders, we looked up the SNP in published results from large GWAS of psychiatric disorders by the PGC (cross-disorder (22), BD (49), MDD (50), and schizophrenia (51)). Here, the cross-disorder GWAS showed the strongest effect, albeit not significant after correction for multiple testing (OR=1.035, unadjusted one-sided *p*=0.048; Supplementary Table S9). Next, we conducted genome-wide comparisons: Using LD score regression, we found no significant genetic correlation between the CCS GWAS and the four psychiatric GWAS (Table 2 and Supplementary Table S10). Further, rank-rank hypergeometric overlap tests showed no significant overlap of SNPs ranked by their association strength (Table 2, Supplementary Table S11, and Supplementary Fig. S5). In binomial sign tests, CCS-associated variants did not show the opposite effect direction in the psychiatric disorder GWAS more often than expected by chance (Table 2 and Supplementary Table S12).

**Table 2:** Comparisons of the component of the common neurobiological substrate (CCS) and the CCS genetic architecture with psychiatric disorders. Details on the four training genome-wide association studies (GWAS) datasets are provided in the Methods section. **LDSC:** Linkage disequilibrium score regression using genome-wide summary statistics (Supplementary Table S10); *r_g_* = genetic correlation. **RRHO:** Rank-rank hypergeometric overlap test showing the relative overlap of genome-wide summary statistics (Supplementary Table S11). **Sign tests:** One-sided binomial sign tests for CCS GWAS *p*-value thresholds *p*<0.05 and *p*<1×10^−5^ and the corresponding probability of success (Supplementary Table S12). **PGS:** Association of polygenic scores with the CCS; *pT* = training GWAS data *p*-value threshold; effect size = linear regression effect size at the *pT* showing the strongest support for an association (see Supplementary Table S13 for results of all ten thresholds); *p*-value: one-sided *p*-value not-corrected for multiple testing. The significance level adjusted for multiple testing was α=0.05/(10×4)=0.00125.

| LD score regression (LDSC) |  |  |
| --- | --- | --- |
| GWAS comparison | $r_g$ | $p$-value |
| Psychiatric Cross-Disorder | 0.0005 | 0.99 |
| Bipolar Disorder | 0.17 | 0.08 |
| Major Depression | -0.03 | 0.75 |
| Schizophrenia | 0.08 | 0.38 |

| Rank-rank hypergeometric overlap (RRHO) |  |  |
| --- | --- | --- |
| GWAS comparison | Overlap | $p$-value |
| Psychiatric Cross-Disorder | 0.29 | 0.53 |
| Bipolar Disorder | 0.21 | 0.06 |
| Major Depression | 0.04 | 0.18 |
| Schizophrenia | 0.02 | 0.15 |

| Binomial sign tests ($p < 0.05$) | | |
| --- | --- | --- |
| GWAS comparison | Probability | $p$-value |
| Psychiatric Cross-Disorder | 0.50 | 0.77 |
| Bipolar Disorder | 0.50 | 0.67 |
| Major Depression | 0.50 | 0.82 |
| Schizophrenia | 0.50 | 0.64 |

| Binomial sign tests ($p < 1 \times 10^{-5}$) | | |
| --- | --- | --- |
| GWAS comparison | Probability | $p$-value |
| Psychiatric Cross-Disorder | 0.33 | 0.93 |
| Bipolar Disorder | 0.54 | 0.50 |
| Major Depression | 0.33 | 0.93 |
| Schizophrenia | 0.54 | 0.50 |

**Table 2:** Comparisons of the component of the common neurobiological substrate (CCS) and the CCS genetic architecture with psychiatric disorders.
| Polygenic scores (PGS) |  |  |  |
| --- | --- | --- | --- |
| Training GWAS | Effect size | $p$-value | $p_T$ |
| Psychiatric Cross-Disorder | -0.78 | 0.30 | $5 \times 10^{-8}$ |
| Bipolar Disorder | -0.64 | 0.05 | $1 \times 10^{-7}$ |
| Major Depression | -5.01 | 0.31 | $1 \times 10^{-2}$ |
| Schizophrenia | -0.58 | 0.24 | $1 \times 10^{-7}$ |

### Analysis of polygenic scores

Next, we calculated PGSs based on the four PGC GWAS (psychiatric cross-disorder, MDD, BD, schizophrenia) as training data and analyzed associations of these disease-associated PGSs with the CCS in our population cohorts. None of the PGSs were associated with the CCS after correction for multiple testing (Table 2, Supplementary Table S13, and Supplementary Fig. S6).

Last, we inverted the direction of the approach and built a PGS based on our CCS GWAS as training data, using ten different *p*-value thresholds, and compared it between patients and controls from four clinical diagnoses (MDD, BD, schizoaffective disorder, and schizophrenia) as available from three patient/control cohorts (BiDirect, MPIP, FOR2107). We expected the CCS PGS to be lower in psychiatric patients. No consistent results were observed regarding the expected direction of the patient/control comparisons and a specific threshold, and no single effect proved robust to multiple testing correction (Supplementary Table S14).

### Analyses of age-dependent effects

In an imaging meta-analysis of our three MDD/control cohorts (BiDirect, MPIP, FOR2107), we confirmed that the CCS was reduced in MDD patients compared to controls (*p*=1.3×10^−7^; Fig. 3A and Supplementary Table S15). In the transdiagnostic FOR2107 cohort, the median CCS showed a stepwise decrease along the affective-psychosis axis (controls: median=0.18; MDD: median=-0.010, comparison to controls: *p*=3.9×10^−3^; BD: median=-0.35, *p*=2.8×10^−5^; schizoaffective disorder: median=-1.13, *p*=2.6×10^−8^; schizophrenia: median=-0.58, *p*=6.6×10^−10^; combined analysis of all four diagnostic groups in FOR2107: *p*=1.5×10^−7^; Fig. 3A, Supplementary Table S15, and Supplementary Fig. S7). This finding strongly affirmed the results of Goodkind *et al*. (1).

**Figure 3:**
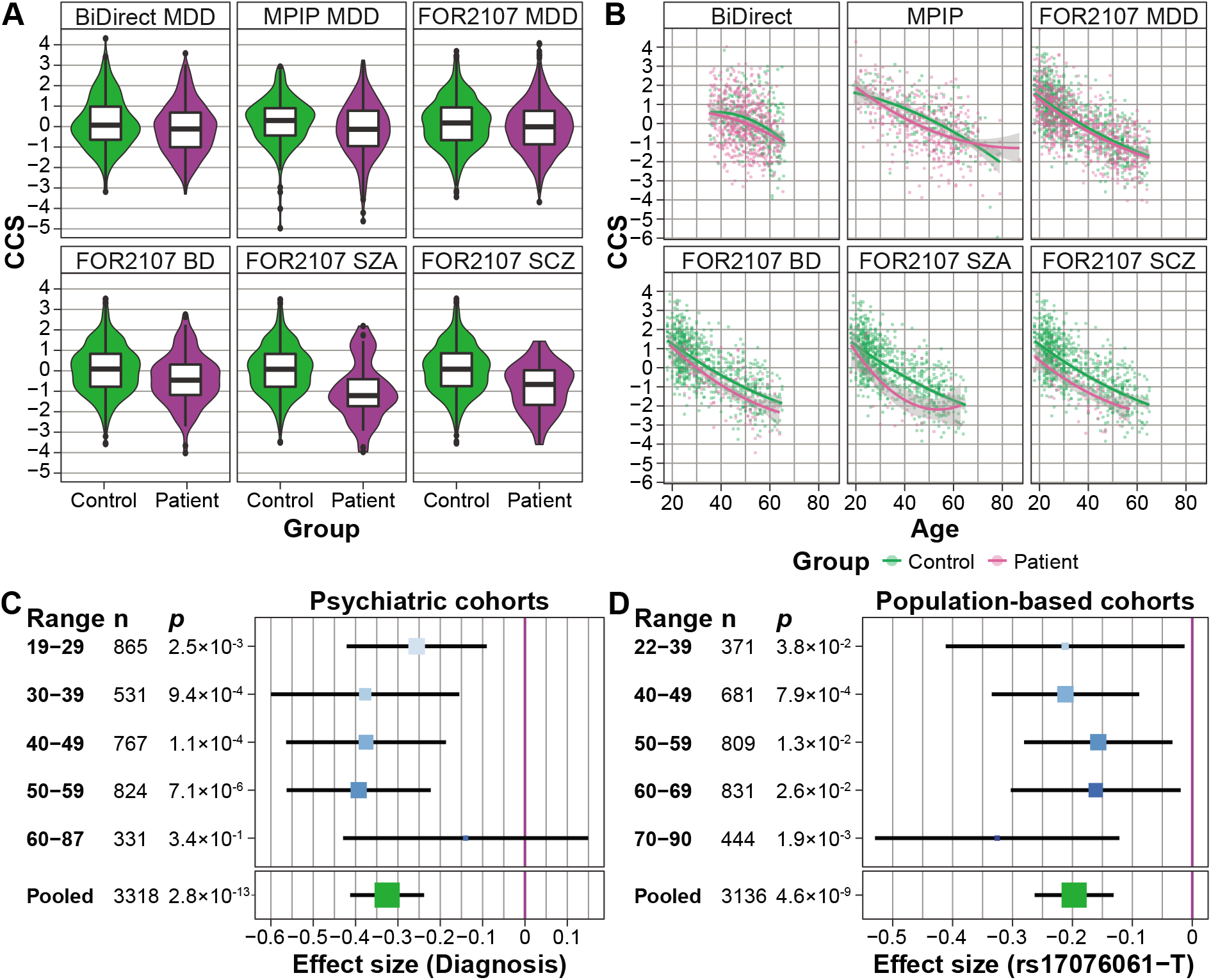
Analyses of age-by-diagnosis and age-by-SNP effects on the component of the common neurobiological substrate (CCS). **A:** A significant smaller CCS was observed in MDD (BiDirect, MPIP, FOR2107), BD (FOR2107), SZA (FOR2107), and SCZ (FOR2107) patients, affirming the transdiagnostic finding by Goodkind *et al*. (1); Supplementary Table S15). **B:** Age^2^ trajectories of the patient/control groups plotted for each cohort. A non-linear, quadratic age dependency was observed in MDD (pooled MPIP, BiDirect, FOR2107), but no other diagnostic group. Data points represent the CCS after residualization against all covariates except for age and age^2^ (separate fit for patients and controls; Supplementary Table S15). **C:** Age-stratified analyses of the association between diagnosis and the CCS using five age groups in the combined patient/control cohorts (with cohort as a covariate). No significant heterogeneity was observed. Size and color of the effect sizes per bin are proportional to the sample size. **D:** Age-stratified analyses of the association between the top SNP (rs17076061) and the CCS using five age groups in the combined five population-based GWAS cohorts (with cohort modelled as a covariate). No significant heterogeneity between the age groups was observed. Size and color of the effect sizes per bin are proportional to the sample size.

In these analyses, we noticed a possible influence of age on the association between the patient/control status and the CCS. When adding a linear and quadratic age interaction term to the MDD regression models, the linear interaction term was not significant (*p*=0.72). However, the age^2^-by-diagnosis interaction term was significant (*p*=0.014), pointing to a possible non-linear age dependency in MDD. No such effect was detected in the other diagnostic groups (Fig. 3B and Supplementary Table S15). To explore non-linear age dependencies in a complementary approach, we stratified all patient/control cohorts into five non-overlapping age groups (Fig. 3C). Heterogeneity in a meta-analysis of the CCS associations stratified by age would have indicated strong non-linear effects of age on the CCS. However, we detected no significant heterogeneity between the age groups (Q=3.21, *p*=0.52; Fig. 3C and Supplementary Table S16).

When adding interaction terms to the model, neither the age-by-SNP (*p*=0.48) nor the age^2^-by-SNP (*p*=0.50) interaction became significant in the meta-analysis of the five GWAS population cohorts, while the main SNP effect remained stable (Supplementary Table S17). Similarly, when stratifying the analysis by age groups, the SNP main effect size varied, yet without significant heterogeneity (Q=2.25, *p*=0.69; Fig. 3D and Supplementary Table S17).

To investigate whether our specific implementation of the global brain size correction influenced the association results, we switched from *non-linear only* Jacobian modulation of the GM probability maps to full Jacobian modulation, with the total intracranial volume entered as an explicit volumetric covariate. Our association results remained stable, independent of the correction method used (Supplementary Methods and Supplementary Table S15).

### Comparison of the genetic architecture of the CCS with the genetics of aging traits

To further explore whether genetic variants associated with the CCS might influence aging-related processes, we compared our CCS GWAS results with GWAS for epigenetic accelerated aging (54) and longevity (53). The common substrate regions represent the salience network, which is specifically prone to neurodegeneration in bvFTD (12, 62), a subtype of frontotemporal dementia with severe executive disturbances and personality changes. Therefore, we also analyzed a possible overlap with GWAS results for bvFTD (52). SNP rs17076061 showed no significant association in any of these GWAS (Supplementary Table S9). Single findings for longevity and epigenetic accelerated aging were nominally significant in PGS analyses and sign tests. However, overall, no significant genetic overlap with any of these GWAS was found with LD score regression (Supplementary Table S10), rank-rank hypergeometric overlap tests (Supplementary Fig. S8 and Supplementary Table S11), sign tests (Supplementary Table S12), or PGS analyses (Supplementary Fig. S6 and Supplementary Table S13) after correction for multiple testing.

## Discussion

In the present study, we investigated the genetic architecture of an MRI-based volumetric marker that has previously been identified as a common neurobiological substrate for major psychiatric disorders (1), mapping to areas of the salience network. As the primary analysis, we conducted a population-based GWAS on this substrate that was calculated from the original three-region substrate using dimensional reduction by PCA. Thereby, we generated the CCS, a construct that simplified our genetic analyses while retaining a large fraction of the phenotypic variance. In secondary analyses, we studied the relationship between the CCS and risk for psychiatric disease as well as age-by-SNP and age-by-diagnosis effects on the CCS. Overall, our study produced three main findings:

First, the minor allele T of the intergenic SNP rs17076061 was associated with a decreased CCS at genome-wide significance and replicated. The association signal from the CCS was stronger than those from the three separate regions indicating that our approach stabilized the CCS association by reducing the statistical noise. The SNP maps directly to an evolutionarily constrained element in mammals (63), supporting a regulatory role of the variant. The locus on chromosome 5q35.2 harbors several predicted, uncharacterized long intergenic non-coding RNAs and two protein-coding genes expressed in the brain with either psychiatric or neuroprotective functions (64–67). The latter genes are “biorientation of chromosomes in cell division 1” (*BOD1*) and “stanniocalcin 2” (*STC2*), located 75 kbp downstream and 202 kbp upstream of rs17076061, respectively.

The SNP is part of a significant expression quantitative trait locus (eQTL) for *STC2* in pancreatic tissue (*p*=3.6×10^−8^). However, this eQTL was not significant in the anterior cingulate cortex (*p*=0.06), and the anterior insula was not available in GTEx v8 (68). Notably, the sample size for the ACC was half of that for the pancreas, decreasing the statistical power. In neurons, rs17076061 thus likely influences the expression of *STC2*, which expresses a secreted glycoprotein with a possible auto- or paracrine function. In the regulation of apoptosis, the unfolded protein response promotes the expression of the potentially neuroprotective STC2 in neuronal cells (66, 67).

Our second main finding is that the neurodevelopmental pathway “SEMA3A-Plexin repulsion signaling by inhibiting Integrin adhesion” was significantly associated with the CCS. Semaphorin-3A (SEMA3A) is a chemorepellent mediating axon guidance and a chemoattractant for dendrite growth, whereas plexins are the signal-transducing subunits of the Semaphorin-3A receptor. Semaphorin-3A and Plexin-A2 are associated with different psychiatric disorders (69–72): *Plexin-A2* is associated with schizophrenia, anxiety, and MDD (72, 73), while *Semaphorin-3A* is upregulated in the brain of schizophrenia patients and has been suggested to contribute to the synaptic pathology of the disorder (70). Furthermore, Semaphorin-3A may contribute to neurodegeneration in Alzheimer’s disease (71), and the pathway is important for neuronal regeneration after brain trauma (74).

A third set of analyses focused on the question whether our approach – correlating a disease-associated structural brain phenotype with population-based genomic variation – would lead to the detection of genetic variants relevant for psychiatric disorders. Here, we found a discrepancy between detecting a genome-wide significant SNP (rs17076061) on the one hand, while not detecting an association between this SNP and major psychiatric diagnoses (MDD, BD, and schizophrenia) on the other hand. This finding obviously contradicts the latent expectation that the CCS could represent a ‘risk endophenotype’ that exhibits a substantial heritability of 50% in the studied population. Although our top SNP explained only a small fraction of the CCS variance (*R^2^*=1.2%, sample size-weighted mean across three cohorts), there still remains a disconnection between this finding and the lack of an observed psychiatric risk conveyed by the SNP.

One explanation for this observation is the low correlation between the CCS and psychiatric diagnoses: Goodkind *et al*. (1) used the revised activation likelihood estimation (ALE) meta-analysis framework to test for a spatial convergence of morphometric patient/control differences and found the three-region substrate. However, ALE does not process effect sizes from the original studies which impeded a comparison with our results. We thus analyzed patient/control cohorts of the affective-psychosis spectrum to assess the CCS variance explained by the diagnostic status, ranging from 1.0% for MDD to 4.2% for schizophrenia (*R^2^*). Therefore, in a model that attributes disease risk to the presence of a smaller CCS (less GM), we expect the risk effect mediated by a single SNP to be very low. Compatible with this model, the association of rs1707601 with disease risk was only nominally significant in the large and most recent cross-disorder study by the PGC (26,432 patients and 49,926 controls; (22)). Evidence from large consortium studies showed that psychiatric disease-specific PGSs explain only a small fraction of the disease phenotype (19). This, along with the low disease/CCS correlation, may explain our observation that PGS calculated from published GWAS were not associated with the CCS in our population-based cohorts.

The polygenic nature of both the CCS and risk for psychiatric disease demanded more detailed comparisons between association signals from the CCS GWAS and GWAS of major psychiatric disorders applying complementary statistical approaches (LD score regression, RRHO, binominal sign tests). Our results suggest that no such genetic overlap exists, adding our study to a line of similar previous reports: Large studies on MDD and schizophrenia, for example, found only weak or no relationship between the genetic architecture of these diagnoses and regional brain volumes (2, 55, 75–77). Conversely, a meta-analysis of genetic factors influencing subcortical volumes in about 40,000 individuals identified a genetic correlation between nucleus accumbens and caudate nucleus for BD, but not for schizophrenia (75). One may speculate that differences between disease-predisposing (‘causal’) brain changes and secondary (‘epiphenomenological’) brain changes (due to substance use or other comorbidities) could play a role for this heterogeneity. Methodologically, the analyses of genetic overlap, as conducted by us and others (61, 75), investigated genome-wide similarities between GWAS. If only some variants showed a joint association or different loci exhibited mixed effect directions, these methods could fail to detect similarities. Similarly, our polygenic scores for a larger CCS were not lower in psychiatric patients diagnosed with MDD, BD, schizoaffective disorder, or schizophrenia. This finding supports that the standard approach of PGS, which only accounts for common *additive* effects, does not adequately capture epistatic gene-by-gene or gene-by-environment effects that influence complex traits and even more, disease risk. Future studies are warranted to explore such relationships based on models that allow for non-additive, particularly interactive effects (78).

Another possible explanation for the dissociation between our genetic findings and disease risk is that other pre-morbid environmental influences, such as the prenatal environment or early life adversity, were not addressed in our study. Such influences could aggravate a morphological risk pattern without being directly reflected in genetic associations. Well-documented examples for these influences are specific correlations between early childhood adversity and salience network dysfunction or GM loss (79–81). In this line of thinking, undetected environmental factors may have shaped the CCS beyond genetic effects in our population cohorts. It is evident that only longitudinal studies of patients and controls can disentangle this challenging question, particularly as longitudinal brain changes themselves show a significant heritable component (82).

In our attempt to understand the function of the top SNP from our GWAS (rs17076061), we considered that aspects of pathological aging (accelerated aging) could play a role. In this regard, reports on different structural brain markers suggest that several major psychiatric diseases are associated with accelerated aging, with different effect sizes and different regional patterns (6, 83). The salience network, in particular, is involved in an accelerated cognitive decline during aging (84). Beyond a cross-sectional replication of small but robust CCS differences between patients and controls, we recognized that the CSS could harbor non-linear age-by-diagnosis interactions in MDD. In fact, the SNP effect proved robust against the inclusion of age-interaction terms, without significant heterogeneity when analyzed in age-binned subgroups. Both results suggest that rs17076061 may have a stable effect on the CSS over the adult lifespan. However, we could not entirely exclude the influence of higher-order non-linear deviations which we could not analyze in the present study. Concordant with this observation, we did not find genetic overlaps between our GWAS and GWAS of longevity (representing an extreme form of healthy aging), or bvFTD (representing an extreme form of salience network degeneration). To clarify the relationship between the CCS and a possibly accelerated salience network aging in psychiatric disease, larger patients/control cohorts are required that allow triple-interaction analyses (genetics, disease status, CSS). Our study has several limitations. First, more comprehensive investigations of age dependencies would have been possible from more homogeneous age distributions in the population cohorts. Still, our main goal demanded to assemble large samples, given the expected small effects of common variants. Second, environmental factors such as childhood adversity were either not available or acquired with heterogeneous instruments in the population cohorts, preventing an inclusion of this dimension as an important source of variance, or interaction factor. Third, the operationalization of the CCS followed the specific result map of Good-kind *etal*. (1), which is a sparse representation of the salience network. Data-driven definitions, e.g. through structural covariance as exemplified before (85), may capture a larger portion of the volumetric variance of the salience network (17, 18).

In conclusion, we detected a replicable, genome-wide significant association of a common variant (rs17076061) with GM areas that represent hubs of the salience network in adult individuals from the general population. The genetic architecture of this network was not correlated with genetic risk for major psychiatric disorders. Future gene-by-environment interaction and functional imaging analyses may enable us to understand how salience-network structure translates to psychiatric disease risk.

GWAS summary statistics are available at: https://kg.ebrains.eu/search/instances/Dataset/40a998fb-9483-42ad-b46b-2f8d0bc5aa3e

The genotype and MRI data can be requested from the individual cohorts. R code for the statistical analyses is available on request.

## Acknowledgments

The study was supported by the German Federal Ministry of Education and Research (BMBF), through the Integrated Network IntegraMent, under the auspices of the e:Med programme (grants 01ZX1314A to MMN and SCi, 01ZX1614J to BMM), by the German Research Foundation (DFG; grant FOR2107: NO246/10-1 to MMN, MU1315/8-2 to BMM), by the European Union’s Horizon 2020 Research and Innovation Programme (grants 785907 (HBP SGA2) to SCi, SCa, and KA and 826421 (VirtualBrainCloud) to SBE), and by the Swiss National Science Foundation (SNSF; grant 156791 to SCi). TFMA was supported by the BMBF through the DIFUTURE consortium of the Medical Informatics Initiative Germany (grant 01ZZ1804A) and by the European Union’s Horizon 2020 Research and Innovation Programme (grant MultipleMS, EU RIA 733161). SBE, KA and FH were supported by the Helmholtz Portfolio Theme Supercomputing and Modeling for the Human Brain, SCa receives funding from the Initiative and Networking Fund of the Helmholtz Association and HM from the BMBF (01ER1205); MMN is a member of the DFG-funded cluster of excellence ImmunoSensation. The BiDirect study is supported by grants of the BMBF to the University of Münster (01ER0816 and 01ER1506). SHIP is part of the Community Medicine Research net of the University of Greifswald, Germany, which is funded by the BMBF (01ZZ9603, 01ZZ0103, and 01ZZ0403), the Ministry of Cultural Affairs and the Social Ministry of the Federal State of Mecklenburg-West Pomerania, and the network ‘Greifswald Approach to Individualized Medicine (GANI_MED)’, funded by the BMBF (03IS2061A). Whole-body MR imaging was supported by a joint grant from Siemens Healthineers, Erlangen, Germany and the Federal State of Mecklenburg West Pomerania. Genome-wide data were supported by the BMBF (03ZIK012). The University of Greifswald is a member of the Caché Campus program of the InterSystems GmbH.

We thank the Heinz Nixdorf Foundation, Germany, for the generous support of the Heinz Nixdorf Recall Study. We thank the respective working groups of the Psychiatric Genomics Consortium (https://www.med.unc.edu/pgc/) for making the summary statistics of their GWAS available.

This work is part of the German multicenter consortium “Neurobiology of Affective Disorders. A translational perspective on brain structure and function^1^’, funded by the DFG (Forschungsgruppe / Research Unit FOR2107). The FOR2107 study was funded by the DFG: grants KI 588/14-1, KI 588/14-2 to TK; DA 1151/5-1, DA 1151/5-2 to UD; NE 2254/1-2 to IN; NO 246/10-1, NO 246/10-2 to MMN; MU1315/8-2 to BMM; extended FOR2107 acknowledgments are available in the Supplement.

We thank the International FTD-Genomics Consortium (IFGC; https://ifgc-site.wordpress.com/) for sharing summary statistics for the bvFTD subgroup. The research group’s affiliations and funding sources can be found in the Supplementary Material. We thank the-research participants and employees of 23andMe, Inc. for their contribution to the MDD meta-analysis published in (52).

