## Supplementary Material for "Genetic factors influencing a neurobiological substrate for psychiatric disorders"

*Table of Contents*


**Supplementary Methods** 3

Image preprocessing and extraction of volumes 3

Principal component analysis (PCA) 4

Genetic quality control (QC) and imputation 4

Sequence of genotype QC 5

Imputation of genotype data 5

Calculation of MDS ancestry components 5

Estimation of the SNP heritability 6

GWAS 6

Generation and analysis of polygenic scores 6

Binomial sign tests 7

**Supplementary Tables** 8

**Table S1:** Demographic data of the samples used in the GWAS. 8

**Table S2:** Demographic data of the psychiatric patient/control samples. 8

**Table S3:** Summary of the genotype QC per dataset. 9

**Table S4:** Median genomic inflation λ. 10

**Table S5:** Full test statistics of genome-wide significant variants. 10

**Table S6:** Pairwise linkage disequilibrium (LD) of the 12 top-associated SNPs. 10

**Table S7:** MAGMA pathway analysis. 10

**Table S8:** MAGENTA pathway analysis. 10

**Table S9:** Lookup of rs17076061 in published GWAS. 10

**Table S10:** Results from LD score regression. 10

**Table S11:** Results from rank-rank hypergeometric overlap tests. 10

**Table S12:** Results from the binomial sign tests. 11

**Table S13:** Test statistics for analyses of polygenic scores based on published GWAS. 11

**Table S14:** Test statistics for the analysis of the CCS GWAS PGS in the psychiatric patient/control cohorts. 11

**Table S15:** Association results of psychiatric diagnoses with the CCS. 12

**Table S16:** Association results stratified by age group. 13

**Table S17:** SNP-by-Age effects. 13

**Supplementary Figures** **14**

**Figure S1:** Population substructure of the merged genotype data of all GWAS samples. 14

**Figure S2:** Quantile-quantile plots of GWAS *p*-values. 15

**Figure S3:** Manhattan plot of GWAS *p*-values on the meta-analysis of discovery and replication cohorts. 16

**Figure S4:** Box-whisker plot of the CCS per genotype of variant rs17076061. 17

**Figure S5:** Rank-rank hypergeometric overlap maps. 18

**Figure S6:** Summary of GWAS-based PGS analyses. 19

**Figure S7:** Box-whisker plot of the CCS per diagnosis group in the FOR2107 cohort sorted by the median CCS. 25

**Figure S8:** Rank-rank hypergeometric overlap maps. 26

**IFGC members and affiliations** **27**

**IFGC acknowledgments** **34**

**FOR2107 acknowledgments** **37**

**MPIP acknowledgments** **38**

### Supplementary Methods

#### Image preprocessing and extraction of volumes

A well-established preprocessing pipeline for voxel-based morphometry (VBM) using diffeomorphic anatomical registration through exponentiated lie algebra (DARTEL) for optimal inter-individual image registration, the VBM8 toolbox for statistical parametric mapping 8 (SPM8) was applied to high-resolution T1-weighted images from individuals of all cohorts. For all analyses, gray matter (GM) 1.5×1.5×1.5 $mm^3$ maps with non-linear only (NLO) Jacobian modulation were created using the VBM8 toolbox. More details of the pipeline are described elsewhere: https://www.fil.ion.ucl.ac.uk/spm/ and http://dbm.neuro.uni-jena.de/vbm/download/
For the extraction of the three spatially disjunct regional GM volumes, binarized versions of the joint result areas of the VBM-based meta-analysis of affective and non-affective disorders were used as extraction masks (see Figure 2 in the study by Goodkind *et al.*).

For secondary follow-up analyses, GM maps with full Jacobian modulation (FJM, *i.e.*, correcting for linear and non-linear spatial warping) were also generated. NLO-based volumes can be considered as intrinsically corrected for a linear factor closely representing head size (http://dbm.neuro.uni-jena.de/vbm/). For FJM-based volumes, total intracranial volume (TIV) was used as a covariate to correct for head size. Here, TIV was calculated as the sum of FJM-modulated GM, white matter, and cerebrospinal fluid, masked by a canonical intracranial volume mask in MNI space. The main GWAS was strictly performed on NLO-based measures. In secondary analyses, the FJM-based measures were used to probe the robustness of the GWAS results towards influences of different types of global volume correction: first, total GM, second, TIV, and, third, estimated head size. Here, the estimated head size corresponds to the linear stretching factor *g* used in NLO-based Jacobian modulation, which is calculated as $g = v_{NLO} / v_{FJM}$, where *v* is the regional volume. FJM-based volumes were also used to disentangle effects of age from effects of brain size; brain size is, unlike TIV, age-related and thus influences the NLO stretching factor *g*.

#### Principal component analysis (PCA)

Before conducting the PCA, we studied the correlations between the three regional, NLO-corrected GM volumes in a merged dataset of all five GWAS cohorts. We calculated Pearson ($r_{raw}$) and partial Pearson ($r_{raw\text{-}partial}$) correlations between the three raw, uncorrected volumes as well as Pearson ($r_{res}$) and partial Pearson ($r_{res\text{-}partial}$) correlations between the three residuals of the linear models (corrected for sex, age, age$^2$, and handedness – see below):

| **Comparison** | **Uncorrected volumes** | | **Residualized volumes** | |
|---|---|---|---|---|
| | $r_{raw}$ | $r_{raw\text{-}partial}$ | $r_{res}$ | $r_{res\text{-}partial}$ |
| Left *vs.* right anterior insular cortex (AIC) | 0.65 | 0.54 | 0.45 | 0.41 |
| Left AIC *vs.* dorsal anterior cingulate cortex (dACC) | 0.52 | 0.33 | 0.28 | 0.19 |
| Right AIC *vs.* dACC | 0.46 | 0.19 | 0.25 | 0.14 |

Eventually, we corrected the three GM volumes (left AIC, right AIC, dACC) for covariates (see the main Methods and Figure 1) in linear models. We scaled and centered the residuals from the linear models and combined them by PCA using the *R* function *prcomp*.

The relative variance explained by the three principal components in the GWAS cohorts were as follows (including the sample size-weighted mean):

| **Cohort** | **Uncorrected volumes** | | | **Residualized volumes** | | |
|---|---|---|---|---|---|---|
| | **PC1** | **PC2** | **PC3** | **PC1** | **PC2** | **PC3** |
| **1000BRAINS** | 0.58 | 0.25 | 0.17 | 0.53 | 0.27 | 0.20 |
| **CONNECT100** | 0.63 | 0.22 | 0.16 | 0.46 | 0.32 | 0.22 |
| **BiDirect** | 0.59 | 0.23 | 0.18 | 0.55 | 0.25 | 0.20 |
| **SHIP-2** | 0.71 | 0.18 | 0.11 | 0.57 | 0.26 | 0.17 |
| **SHIP-Trend** | 0.72 | 0.17 | 0.11 | 0.57 | 0.26 | 0.17 |
| **Weighted mean** | **0.67** | **0.20** | **0.13** | **0.55** | **0.26** | **0.18** |

The loadings of the first principal component (*i.e.*, the component of the common substrate, CCS) in the different cohorts were as follows:

| | **Left AIC** | **Right AIC** | **dACC** |
|---|---|---|---|
| **1000BRAINS** | 0.6281114 | 0.6002601 | 0.4951403 |
| **CONNECT100** | 0.6879444 | 0.6149882 | 0.3853855 |
| **BiDirect** | 0.6064360 | 0.6014543 | 0.5200847 |
| **SHIP-2** | 0.6310510 | 0.6072448 | 0.4827302 |
| **SHIP-Trend** | 0.6178612 | 0.6226124 | 0.4802098 |
| **MPIP patients/controls** | 0.6174913 | 0.5980982 | 0.5108649 |
| **BiDirect patients/controls** | 0.5991645 | 0.6266227 | 0.4983431 |
| **FOR2107 patients/controls** | 0.6300093 | 0.6354365 | 0.4464401 |

#### Genetic quality control (QC) and imputation

QC of genotype data was conducted for each dataset separately using PLINK v1.90b2m (or higher versions) and *R* (see Table S3).

##### Sequence of genotype QC

1. Removal of SNPs with call rates <98% or a minor allele frequency (MAF) <1%
2. Removal of individuals with genotyping rates <98%
3. Removal of sex mismatches
4. Removal of genetic duplicates
5. Removal of cryptic relatives with pi-hat≥12.5
6. Removal of genetic outliers with a distance from the mean of >4 SD in the first eight multidimensional scaling (MDS) ancestry components
7. Removal of individuals with a deviation of the autosomal or X-chromosomal heterozygosity from the mean >4 SD
8. *Autosomal variants only*: Removal of non-autosomal variants
9. *X chromosome only*: Removal of variants with significantly different MAFs between males and females (using XWAS)
10. Removal of SNPs with call rates <98% or a MAF <1% or Hardy-Weinberg Equilibrium (HWE) test *p*-values $<1\times10^{-6}$
11. Removal of A/T and G/C SNPs
12. Update of variant IDs and positions to the IDs and positions in the 1000 Genomes Phase 1 reference panel
13. Alignment of alleles to the reference panel
14. Removal of duplicated variants and variants not present in the reference panel

For the X chromosome, QC was conducted separately for males and females. Note that the 1000BRAINS CoreExome samples were imputed as part of a larger dataset to improve imputation quality; the relevant individuals were extracted from the dataset after imputation.

##### Imputation of genotype data

Genotypes were aligned to the 1000 Genomes Phase 1 reference panel using SHAPEIT v2 (r790 or higher) and PLINK v1.90b2m (or higher). Pre-phasing (haplotype estimation) was conducted for each chromosome separately using SHAPEIT. Imputation was performed using IMPUTE2 v2.3.1 (or higher) in 5 Mbp chunks with 500 kbp buffers, filtering out variants that are monomorphic in the EUR samples. Chunks with <51 genotyped variants or concordance rates <92 % were fused with neighboring chunks and re-imputed. Variants with a MAF <1% or an INFO metric <0.8 were removed after imputation.

#### Calculation of MDS ancestry components

For the population substructure analysis, pre-imputation genotype data were used, after the QC steps explained above had been applied. Additional variant filtering steps were: removal of variants with a MAF <0.05 or HWE *p*-value $<10^{-3}$; removal of variants mapping to the extended MHC region (chromosome 6, 25-35 Mbp) or to a typical inversion site on chromosome 8 (7-13 Mbp); linkage disequilibrium (LD) pruning (command `--indep-pairwise 200 100 0.2`). Next, the pairwise identity-by-state (IBS) matrix of all individuals was calculated using the

command `--genome` on the filtered genotype data. Multidimensional scaling (MDS) analysis was performed on the IBS matrix using the eigendecomposition-based algorithm in PLINK v1.90b3.38 or higher.
MDS ancestry components calculated for each cohort separately were used as covariates in analyses using genetic data.

#### Estimation of the SNP heritability

Genomic-relatedness-based restricted maximum-likelihood (GREML), as implemented in genome-wide complex trait analysis (GCTA), was used to estimate the SNP heritability $h^2_g$. To this end, the imputed data from all cohorts were converted to best-guess genotypes (`--hard-call-threshold 0.1`) and merged in PLINK. In the merged dataset, only variants present in the HapMap r28 CEU release, with a call rate ≥0.98, and a MAF ≥0.01 were retained. The merged genotype data was LD-pruned using the PLINK command `--indep-pairwise 200 100 0.5`. Subsequently, 137,283 variants were used for the calculation of the genetic relationship matrix in GCTA. The genotyping microarray type and eight MDS ancestry components were used as covariates in the analysis. GREML was conducted using the option `--grm-adj 0`, assuming that causal variants have a similar distribution of allele frequencies as the genotyped variants.

#### GWAS

GWAS of the NLO-based CCS was conducted in PLINK separately per cohort, with eight MDS components as covariates. For 1000BRAINS, the imputation batch was used as an additional covariate. Variants on the X chromosome (non-pseudoautosomal region) were analyzed separately for males and females, followed by combination using the *p*-value-based *SAMPLESIZE* method in METAL to allow for different effect sizes per sex. A two-stage design was implemented, including discovery and replication, with SHIP-Trend used as an independent replication sample. Cohorts were combined in both stages by fixed-effects meta-analysis using METAL, and only variants present in all discovery cohorts were analyzed. LD of SNPs showing an association with $p<5\times10^{-8}$ in the discovery stage was analyzed using the CEU population in LDmatrix. The SNPs that showed the most robust support for an association and, where applicable, with an LD $r^2<0.5$ with more strongly associated variants, were carried forward to the replication stage. Replication criterion was a one-sided *p*-value <0.05/n, where n=2 for the two variants analyzed in the replication cohort. The association statistics of genome-wide significant variants were confirmed in *R* v3.4.3. The residuals of the linear models of genome-wide significant variants were normally distributed in all cohorts (Shapiro-Wilk test).
In each cohort, at least two of the twelve SNPs associated with genome-wide significance were genotyped on microarrays; the remaining ten variants were imputed (imputation quality INFO ≥0.975).

#### Generation and analysis of polygenic scores

For each polygenic score (PGS), the effect sizes of variants below a selected *p*-value threshold, both obtained from either previously published or the present GWAS (training data), were multiplied by the imputed SNP dosage in the test data and then summed to produce a single PGS per threshold. A common set of variants imputed

in all five cohorts was determined. The GWAS training summary statistics and this common set of variants were merged based on chromosome, position, and alleles of each variant. Summary statistics were then clumped in PLINK v1.90b6.4 or higher, based on best-guess genotype data (hard-call threshold 0.3) using the following parameters:
`--clump-kb 500 --clump-r2 0.1 --clump-p1 1 --clump-p2 1`
PGS were then calculated in *R* v.3.3 or higher based on imputed (dosage) data. Test statistics and alleles in the GWAS training data were flipped so that effect sizes were always positive. Thus, the PGS represent weighted, cumulative, additive risk. PGS were scaled to represent the relative risk load (minimum possible cumulative risk load = 0, maximum = 1). For each disorder, ten PGS with different *p*-value thresholds were calculated: $<5\times10^{-8}$, $<1\times10^{-7}$, $<1\times10^{-6}$, $<1\times10^{-5}$, $<1\times10^{-4}$, <0.001, <0.01, <0.05, <0.1, <0.2.
PGS calculated from previously published GWAS were analyzed using linear regression models separately per cohort in *R* v3.4.3, using the regression formula *CCS ~ PGS + covariates*. CCS PGS were analyzed in the case/control cohorts using logistic regression with the model *Diagnosis ~ PGS + covariates*. Eight MDS ancestry components were used as covariates to control for population substructure, and in case of 1000BRAINS, imputation batches. Summary statistics from the cohorts were combined using fixed-effects meta-analysis. Report *p*-values have not been corrected for multiple testing. Significance thresholds were corrected for multiple testing using the Bonferroni method: per GWAS, PGS with ten different *p*-value thresholds were calculated, thus, $\alpha = 0.05/10 = 0.005$. One-sided test statistics were calculated according to the hypothesis that the CCS is negatively correlated to the psychiatric disorder PGS (linear regression analyses of PGS for previous GWAS) or that the CCS PGS is lower in psychiatric patients (logistic regression analyses of the CCS PGS).

#### Rank-rank hypergeometric overlap tests

The rank-rank hypergeometric overlap (RRHO) test was used to compare whether the order of SNPs ranked by their association strength between studies was random. For this analysis, variants were LD-pruned in the EUR subset of the 1000 genomes phase 3 reference panel in PLINK using the command `--indep-pairwise 200 5 0.25`. The RRHO step size *s* was adapted based on the SNP overlap between studies: <50,000 SNPs: *s*=1000, ≥50,000 and ≤150,000 SNPs: *s*=2000, >150,000 SNPs: *s*=3000.

#### Binomial sign tests

Binomial sign test was conducted in *R* using the function *binom.test* to analyze whether SNPs associated with the CCS at either $p<0.05$ or $p<1\times10^{-5}$ showed the opposite direction of effects in psychiatric disorder GWAS more often than expected by chance. The opposite direction was used because the CCS should be smaller in patients with a psychiatric disorder. For this analysis, CCS GWAS meta-analysis summary statistics were LD-clumped in the EUR subset of the 1000 genomes phase 3 reference panel (with MAF≥0.05) in PLINK using the following parameters:
`--clump-kb 500 --clump-r2 0.1 --clump-p1 1 --clump-p2 1`.
LD-independent sign tests were calculated using the lead variant per LD block.

### Supplementary Tables

**Table S1:** Demographic data of the samples used in the GWAS.
The table lists samples after genotypic and phenotypic QC.
Age: mean (SD). Sex: n (%). In 1000BRAINS, CONNECT100, and BiDirect, psychiatric diagnoses or symptoms were collected with self-report questionnaires. In SHIP-2 and SHIP-Trend, diagnoses were assessed using the M-CIDI interview. Dep.: the number of probands with (self-reported) depression or depressive symptomatology. Other: In 1000BRAINS, the number of probands with a self-reported diagnosis of obsessive-compulsive disorder; in BiDirect, the number of probands with a self-reported diagnosis of anxiety or anorexia nervosa; in SHIP-2, the number of probands with a diagnosis of BD. NA: information not available. The observed frequency of psychiatric diagnoses corresponds to the prevalence in the general population and is thus expected in population-based cohorts.

| Cohort | n | Age | Sex (female) | Dep. (%) | Other (%) |
|---|---|---|---|---|---|
| 1000BRAINS | 539 | 67.7 (6.7) | 248 (46.0 %) | 19 (3.5) | 1 (0.2) |
| CONNECT100 | 93 | 47.4 (13.8) | 49 (52.7 %) | 4 (4.3) | NA |
| BiDirect | 589 | 52.8 (8.1) | 301 (51.1 %) | 8 (13.6) | 12 (2.0) |
| SHIP-2 | 1050 | 55.8 (12.7) | 542 (51.6 %) | 137 (13.0) | 4 (0.4) |
| SHIP-Trend | 865 | 50.5 (13.5) | 477 (55.1 %) | 144 (16.6) | NA |
| **Total** | **3136** | **55.6 (12.9)** | **1617 (51.6 %)** | **312 (9.9)** | **17 (0.5)** |

**Table S2:** Demographic data of the psychiatric patient/control samples.
The table lists samples after genotypic and phenotypic QC. MDD: Major Depressive Disorder, BD: Bipolar Disorder, SCZ: Schizophrenia, SZA: Schizoaffective Disorder.
Age: mean (SD). Sex: n (%).
$p_{Age}$: $p$-value of t-tests for differences in age between patients and controls;
$p_{Sex}$: $p$-value of $X^2$-tests for differences in sex between patients and controls.

| Cohort | n | Age | $p_{Age}$ | Sex (female) | $P_{Sex}$ |
|---|---|---|---|---|---|
| BiDirect controls | 311 | 53.1 (7.95) | | 149 (47.9 %) | |
| BiDirect MDD patients | 582 | 49.4 (7.25) | $4.2\times10^{-11}$ | 338 (58.1 %) | $4.6\times10^{-03}$ |
| MPIP patients | 197 | 49.6 (13.6) | | 114 (57.9 %) | |
| MPIP MDD patients | 385 | 48.1 (13.7) | $2.0\times10^{-01}$ | 205 (53.2 %) | $3.3\times10^{-01}$ |
| FOR2107 all controls | 867 | 34.0 (12.7) | | 550 (63.4 %) | |
| FOR2107 MDD controls | 831 | 34.0 (12.5) | | 521 (62.7 %) | |
| FOR2107 MDD patients | 769 | 36.7 (13.3) | $3.3\times10^{-05}$ | 505 (65.7 %) | $2.4\times10^{-01}$ |
| FOR2107 BD controls | 831 | 34.0 (12.5) | | 521 (62.7 %) | |
| FOR2107 BD patients | 127 | 41.4 (12.0) | $1.5\times10^{-09}$ | 69 (54.3 %) | $8.8\times10^{-02}$ |
| FOR2107 SCZ controls | 828 | 34.0 (12.5) | | 519 (62.7 %) | |
| FOR2107 SCZ patients | 72 | 37.2 (10.8) | $1.7\times10^{-02}$ | 30 (41.7 %) | $7.2\times10^{-04}$ |
| FOR2107 SZA controls | 832 | 34.0 (12.5) | | 522 (62.7 %) | |
| FOR2107 SZA patients | 43 | 38.8 (12.8) | $2.1\times10^{-02}$ | 23 (53.5 %) | $2.9\times10^{-01}$ |
| **Total** | **3,353** | **42.0 (13.8)** | | **1,983 (59.1 %)** | |

**Table S3:** Summary of the genotype QC per dataset.
For a detailed description of the QC steps, please see the section Sequence of genotype QC (page 5). Some individuals from the 1000BRAINS cohort were genotyped twice on different microarrays. Hence, the sum of the samples available before QC on the three 1000BRAINS microarrays appears higher in the table below than the total number of individuals in the combined cohort. In cases where individuals were genotyped on several microarrays, the sample with more variants after imputation was selected.

| Dataset / imputation batch | Microarray | Individuals before QC | Individuals after QC | Variants before QC | Variants after QC | Variants after imputation |
|---|---|---|---|---|---|---|
| 1000BRAINS / CONNECT100 *OmniExpress* | Illumina OmniExpress | 533 | 460 | 719,666 | 634,222 | 8,575,622 |
| 1000BRAINS *Omni1-Quad* | Illumina Omni1-Quad | 143 | 123 | 1,134,514 | 744,227 | 8,620,154 |
| 1000BRAINS *CoreExome* | Illumina CoreExome | 52 | 49 * | 538,448 | 246,055 | 7,885,975 |
| 1000BRAINS / CONNECT100 *all three microarrays combined after imputation (no duplicates)* | Combined (summary of the final dataset) | 714 | 632 | *merged after imputation* | *merged after imputation* | 7,570,797 |
| BiDirect GWAS | Illumina PsychArray | 623 | 589 | 588,454 | 262,296 | 8,061,563 |
| SHIP-2 | Affymetrix 6.0 | 1,142 | 1,050 | 900,475 | 524,983 | 8,222,555 |
| SHIP-Trend | Illumina Omni 2.5 | 896 | 865 | 2,450,000 | 1,466,103 | 8,921,041 |
| BiDirect MDD patient/control | Illumina PsychArray | 2,129 | 893 * | 588,454 | 283,900 | 8,599,860 |
| MPIP MDD patient/control *610k* | Illumina Human610-Quad | 225 | 225 | 716,385 | 534,384 | *merged before imputation* |
| MPIP MDD patient/control *550k* | Illumina 550k | 321 | 320 | 522,008 | 502,481 | *merged before imputation* |
| MPIP MDD patient/control *OmniExpress* | Illumina OmniExpress | 135 | 123 | 716,503 | 595.204 | *merged before imputation* |
| MPIP MDD patient/control *all three microarrays combined before imputation* | Combined (summary of the final dataset) | 668 | 582 | 274,863 | 274,863 | 8,470,491 |
| FOR2107 patient/control | Illumina PsychArray | 2,375 | 1886 * | 596,861 | 284,694 | 8,565,139 |

* Imputation was conducted as part of a larger sample and eligible individuals for this study were extracted after imputation.

**Table S4:** Median genomic inflation λ.
The genomic inflation was in the expected range.
For the meta-analyses, λ normalized for 1000 individuals ($\lambda_{1000}$) is also provided.
λ was calculated on the autosomal chromosomes 1-22.

| Analysis | n | λ | $\lambda_{1000}$ |
|---|---|---|---|
| 1000BRAINS | 539 | 1.0075 | |
| CONNECT100 | 93 | 1.0051 | |
| BiDirect | 589 | 1.0005 | |
| SHIP-2 | 1050 | 0.9977 | |
| SHIP-Trend | 865 | 0.9977 | |
| Discovery | 2271 | 1.0240 | 1.0106 |
| Discovery+Replication | 3136 | 1.0283 | 1.0090 |

**Table S5:** Full test statistics of genome-wide significant variants.
BETA = effect size relative to allele 1, SE = standard error, P = *p*-value. R2 = variance explained by the full model (and not by the SNP alone).
This table is provided in the separate Excel file.

**Table S6:** Pairwise linkage disequilibrium (LD) of the 12 top-associated SNPs. Analyzed using the 1000 Genomes CEU population in LDmatrix (https://ldlink.nci.nih.gov/). This table is provided in the separate Excel file.

**Table S7:** MAGMA pathway analysis.
The table shows all nominally significant (P<0.05) Reactome pathways.
P_CORR = *p*-value corrected for multiple testing using the Holm method.
This table is provided in the separate Excel file.

**Table S8:** MAGENTA pathway analysis.
The table shows all nominally significant (either NOMINAL_GSEA_PVAL_95PERC_CUTOFF<0.05 or NOMINAL_GSEA_PVAL_75PERC_CUTOFF<0.05) Reactome pathways.
This table is provided in the separate Excel file.

**Table S9:** Lookup of rs17076061 in published GWAS.
Lookup of the association coefficients from published summary statistics of the psychiatric disorder and aging-related GWAS, as referenced in the main text. For longevity, BETA = positive indicates that no effect size was provided in the GWAS summary statistics; allele T of the SNP is associated with longevity (not significant after correction for multiple testing).
This table is provided in the separate Excel file.

**Table S10:** Results from LD score regression.
This table is provided in the separate Excel file.

**Table S11:** Results from rank-rank hypergeometric overlap tests.
Comparison of CCS-associated variants to variants associated in GWAS for either psychiatric disorders (first part) or aging-related traits (second part).

N = number of variants. Relative overlap = *n* overlapping / *n* SNPs. The step size was adapted according to the *n* SNPs in common between two GWAS. Also see Supplementary Figs. S5 and S8. This table is provided in the separate Excel file.

**Table S12:** Results from the binomial sign tests.
CI = confidence intervals. This table is provided in the separate Excel file.

**Table S13:** Test statistics for analyses of polygenic scores based on published GWAS.
Summary statistics from linear regression analyses to test for an association of PGS with the CCS. The first part of the table shows associations with psychiatric disorder PGS, the second part with aging-related PGS. One-sided test statistics were calculated according to the hypothesis that the CCS is negatively correlated to the psychiatric disorder PGS, *i.e.*, that individuals with a smaller CCS have higher genetic risk load for disorders. Also see Supplementary Fig. S6. bvFTD = behavioral frontotemporal dementia, PC = prefrontal cortex.
This table is provided in the separate Excel file.

**Table S14:** Test statistics for the analysis of the CCS GWAS PGS in the psychiatric patient/control cohorts.
Summary statistics from logistic regression analyses to test for an association of CCS PGS with patient/control status. One-sided test statistics were calculated according to the hypothesis that the CCS PGS is lower in psychiatric patients. MDD: Major Depressive Disorder, BD: Bipolar Disorder, SZA: Schizoaffective Disorder, SCZ: Schizophrenia. This table is provided in the separate Excel file.

**Table S15:** Association results of psychiatric diagnoses with the CCS.
This table shows summary statistics from linear regression analyses and is provided in the separate Excel file.
The upper sub-table shows the analyses of MDD patient/control cohorts. The lower sub-table shows results for the transdiagnostic FOR2107 (FOR) cohort with the following diagnoses: MDD: Major Depressive Disorder, BD: Bipolar Disorder, SCZ: Schizophrenia, SZA: Schizoaffective Disorder, All: all available diagnoses.

*Explanation of the 'trait' column:*
**CCS:** Standard calculation of CCS, based on NLO volumes
1. Extraction of volumes from SPM using non-linear only (NLO) modulation
2. Correction of individual volumes for covariates (age, $age^2$, sex, handedness) in linear regression models
3. Calculation of PCA based on regression residuals
4. CCS association analysis of the first principal component with ancestry components as covariates

**CCS (without residualization):** Calculation of CCS based on native NLO volumes
1. Extraction of volumes from SPM using NLO modulation
2. Calculation of PCA based on the NLO volumes (not corrected for covariates by residualization)
3. CCS association analysis of the first principal component with ancestry components and all other covariates (age, $age^2$, sex, handedness)

**CCS (FJM):** Standard calculation of CCS, based on FJM volumes
1. Extraction of volumes from SPM using full Jacobian (FJM) modulation
2. Correction of individual volumes for covariates (total intracranial volume (ICV), age, $age^2$, sex, handedness) in linear regression models
3. Calculation of PCA based on regression residuals
4. CCS association analysis of the first principal component with ancestry components as covariates

*Explanation of the 'Interaction term' column:*

| | |
|---|---|
| **None:** | No interaction terms were included in the models |
| **Age*diagnosis+Age^2*diagnosis:** | age × diagnosis and $age^2$ × diagnosis interaction terms were included in the regression models after PCA |

*Explanation of the 'Coefficients for' column:*
Here, the term of the model is specified for which the regression coefficients are provided.

**Table S16:** Association results stratified by age group.
The first sub-table shows test statistics for the association of MDD diagnosis with the CCS (MDD cohorts). The second sub-table shows test statistics for the association of any diagnosis with the CCS (all patient/control cohorts). The third sub-table shows test statistics for the association of SNP rs17076061 with the CCS (GWAS cohorts). Individuals were grouped by decade; neighboring groups were merged where <50 individuals per decade. Analyses were conducted on a combined dataset of all cohorts, including a covariate indicating cohort. Q = Cochran's Q from the meta-analysis. This table is provided in the separate Excel file.

**Table S17:** SNP-by-Age effects.
Detailed effects of interactions and VBM modulation on regression coefficients in the association of rs17076061 (SNP) with the CCS. The 'trait' column is organized as explained for Table S15. Furthermore, 'genotyping batch' was included as a covariate for analyses of the 1000BRAINS cohort.
This table shows summary statistics from linear regression analyses and is provided in the separate Excel file.

*Explanation of the 'interaction term' column:*

| | |
|---|---|
| **None:** | No interaction terms were included in the models |
| **age*SNP+age^2*SNP:** | age × SNP and $age^2$ × SNP interaction terms were included in the regression models after PCA. SNP = rs17076061 |

### Supplementary Figures

**Figure S1:** Population substructure of the merged genotype data of all GWAS samples.
Note that the MDS analysis on the merged dataset can differ from the MDS analyses of the individual cohorts (the ancestry components used as covariates in the GWAS).

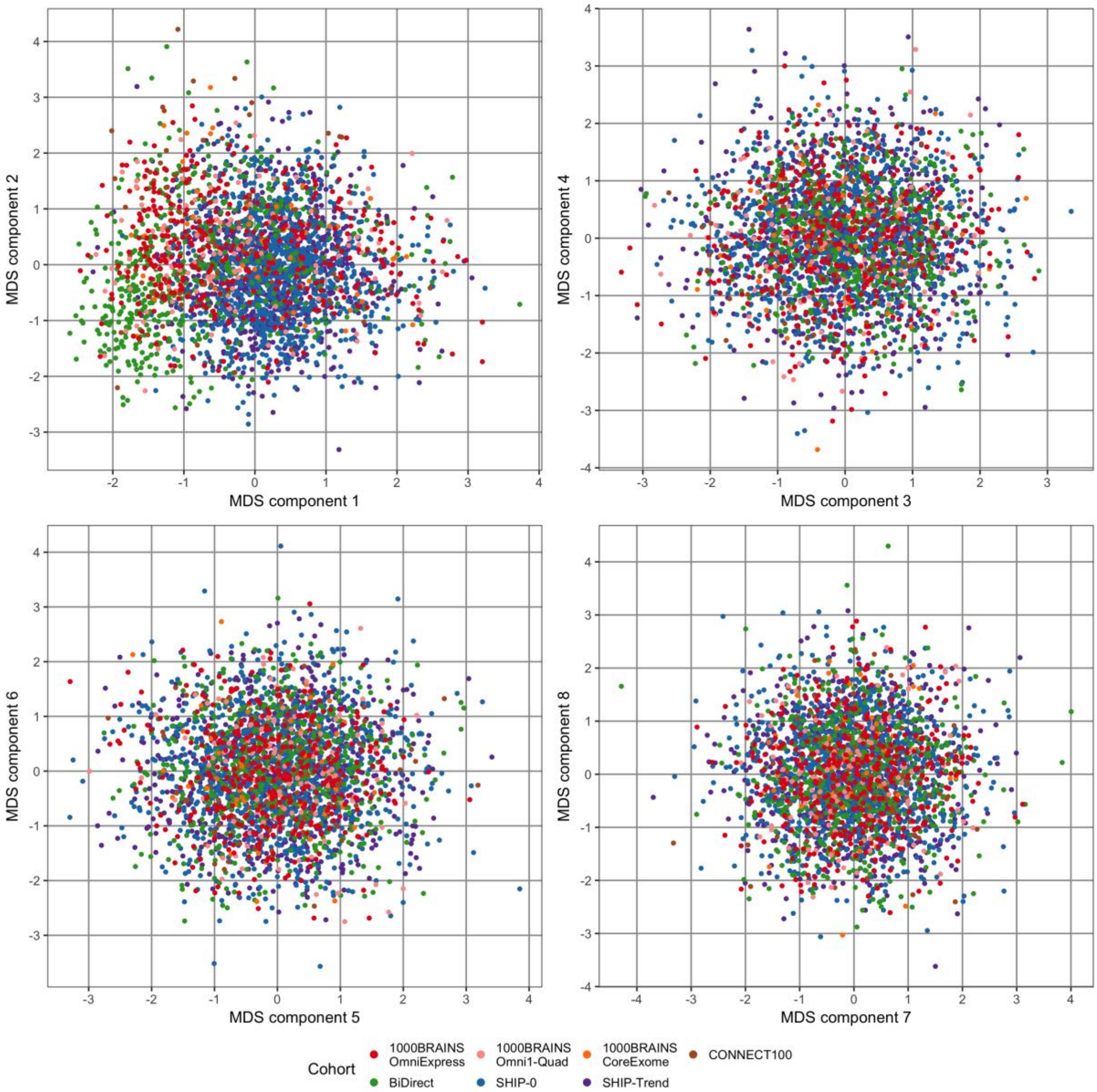

**Figure S2:** Quantile-quantile plots of GWAS *p*-values.
**A:** Quantile-quantile plot for the discovery stage GWAS.

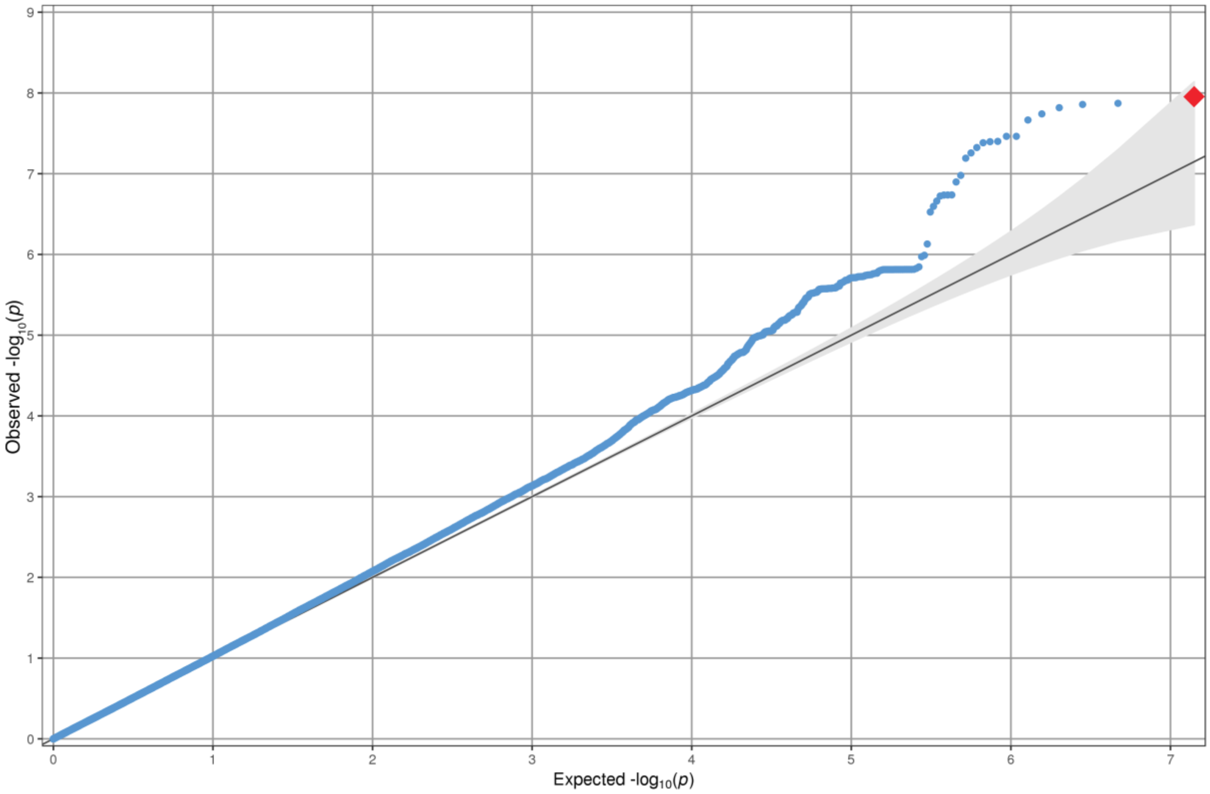


**B:** Quantile-quantile plot for the meta-analysis of discovery and replication cohorts.

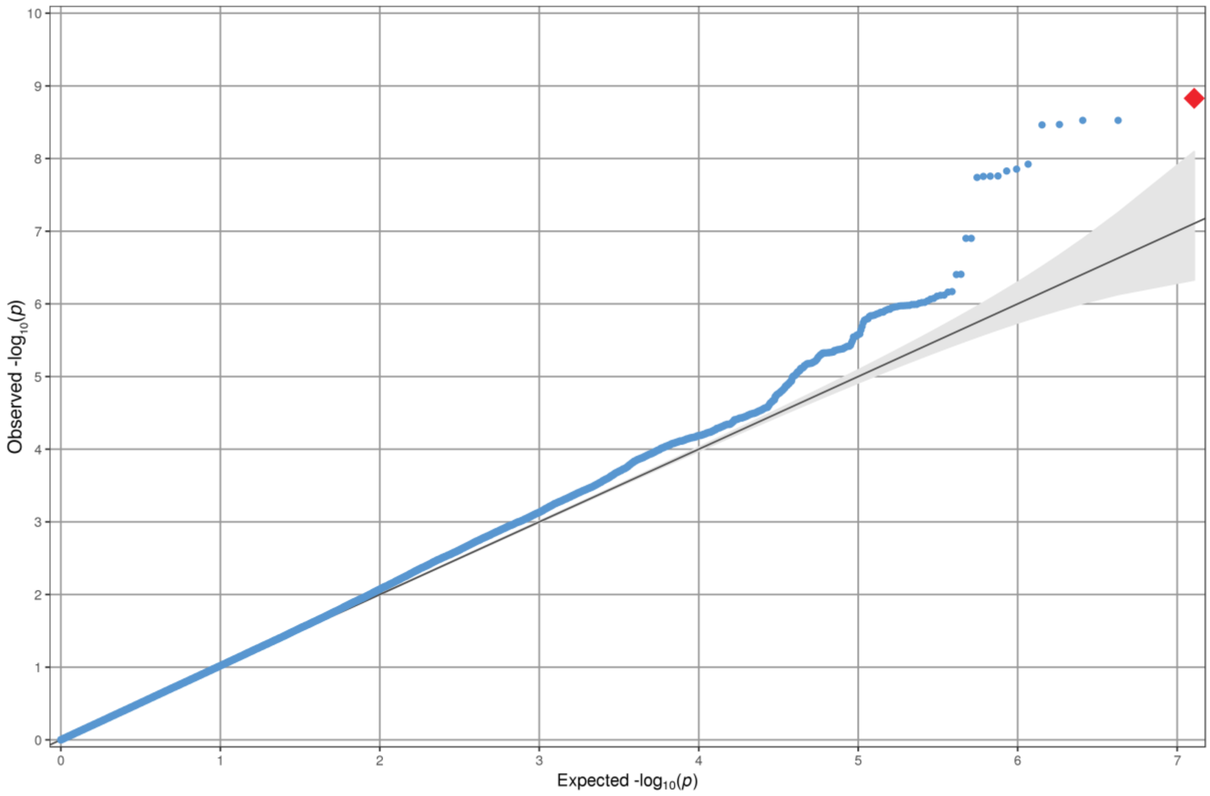

**Figure S3:** Manhattan plot of GWAS *p*-values on the meta-analysis of discovery and replication cohorts.
Red line: genome-wide significance threshold.

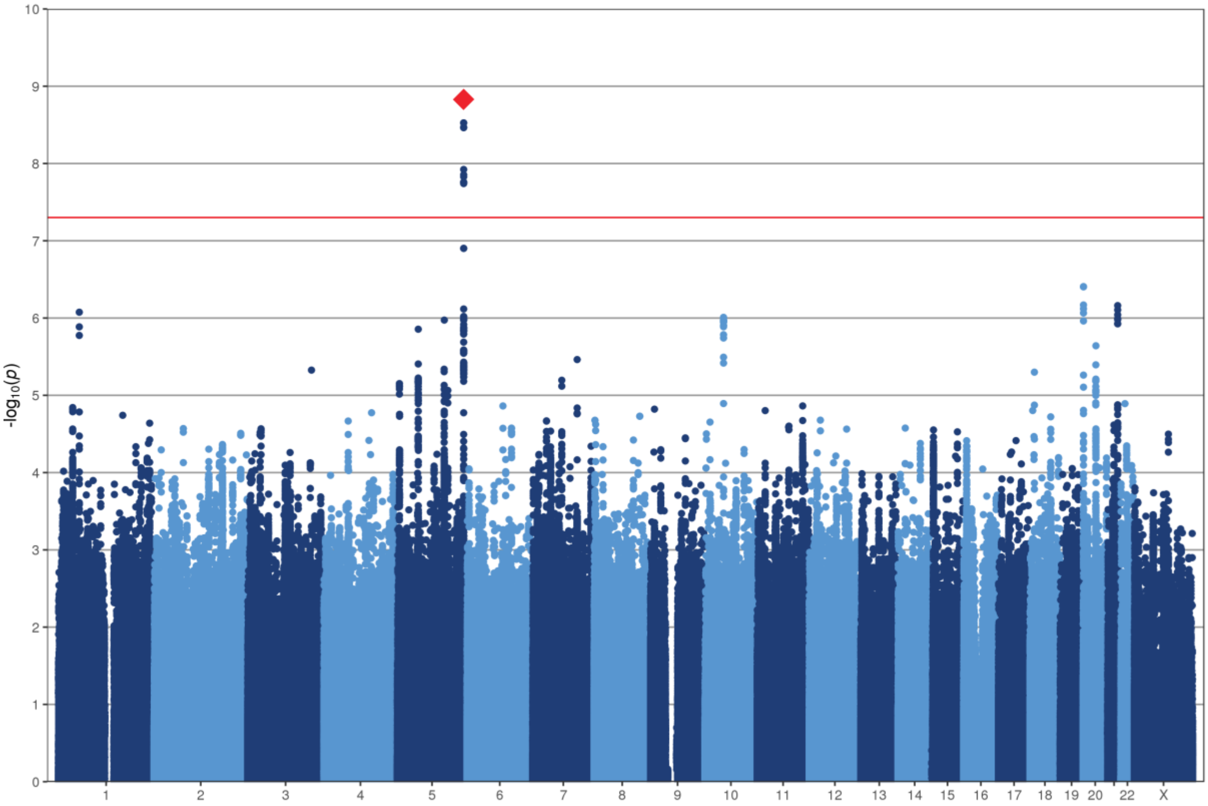

**Figure S4:** Box-whisker plot of the CCS per genotype of variant rs17076061.

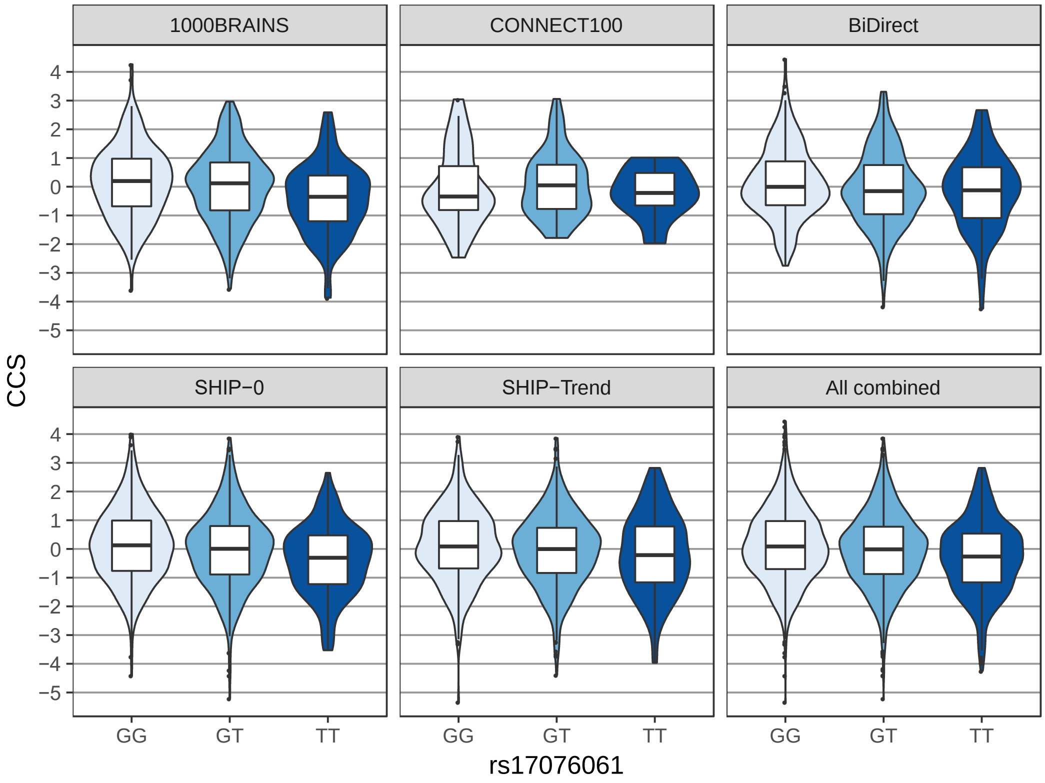

**Figure S5:** Rank-rank hypergeometric overlap maps.
Comparison of CCS-associated and psychiatric disorder GWAS-associated variants. For further details see Table S11. Note that A shows the Cross-Disorder 2019 GWAS.

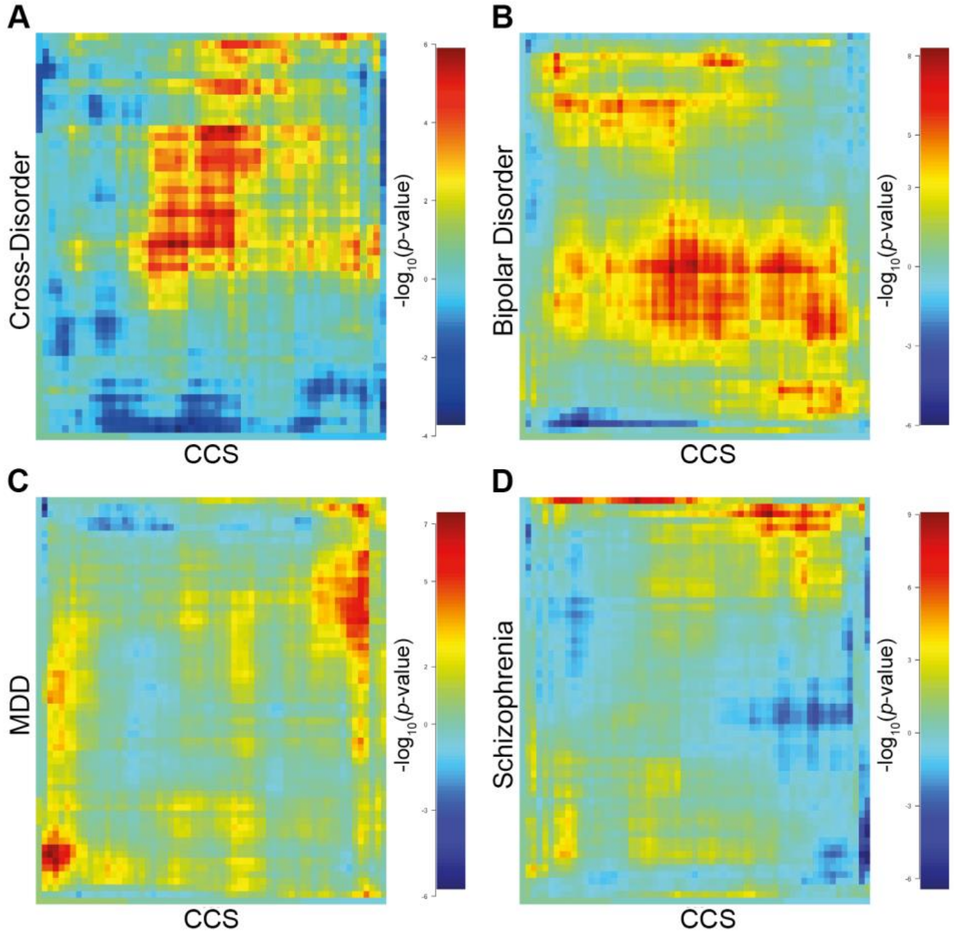

**Figure S6:** Summary of GWAS-based PGS analyses.
Results from linear regression analyses to test for the association of psychiatric disorder and aging-related GWAS PGS with the CCS. The plots show one-sided *p*-values following the hypothesis that individuals with a small CCS have higher disorder or aging-related PGS. The PGS at different training GWAS *p*-value thresholds are shown on the x-axis, and the association strength (-$\log_{10}$ *p*) is shown on the y-axis. Orange dashed line: nominal significance threshold (α=0.05). See Table S13 for further details.

**A:** Analysis of PGS based on the Cross-Disorder 2013 GWAS as training data.

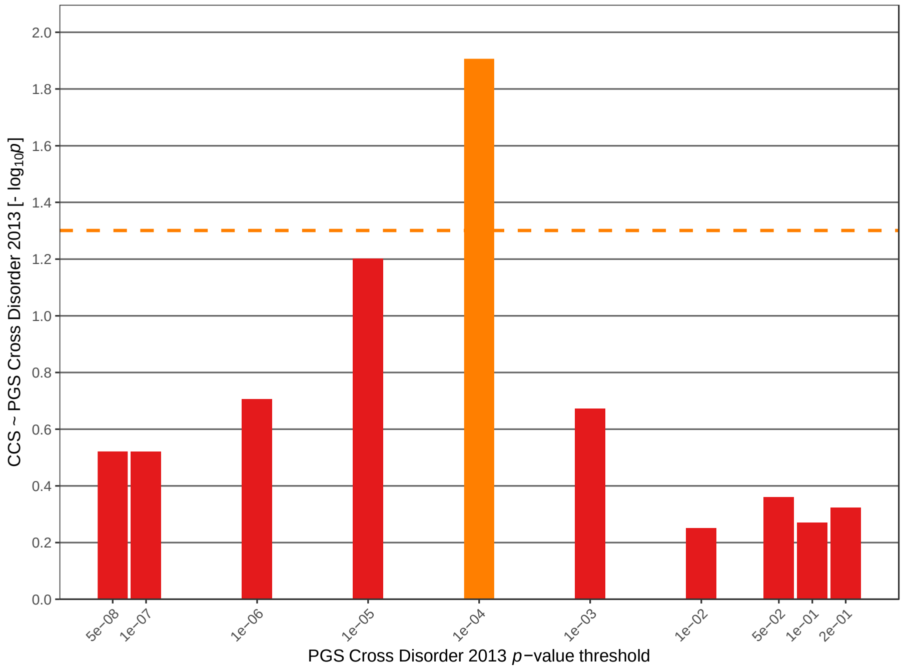


**B:** Analysis of PGS based on the Cross-Disorder 2019 GWAS as training data.

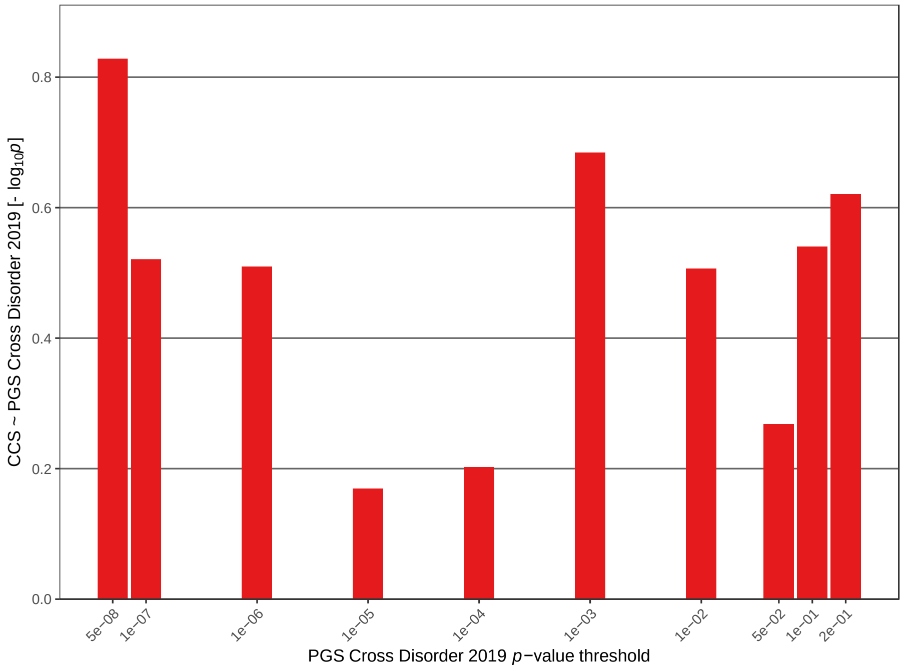


**C:** Analysis of PGS based on the Bipolar Disorder GWAS as training data.

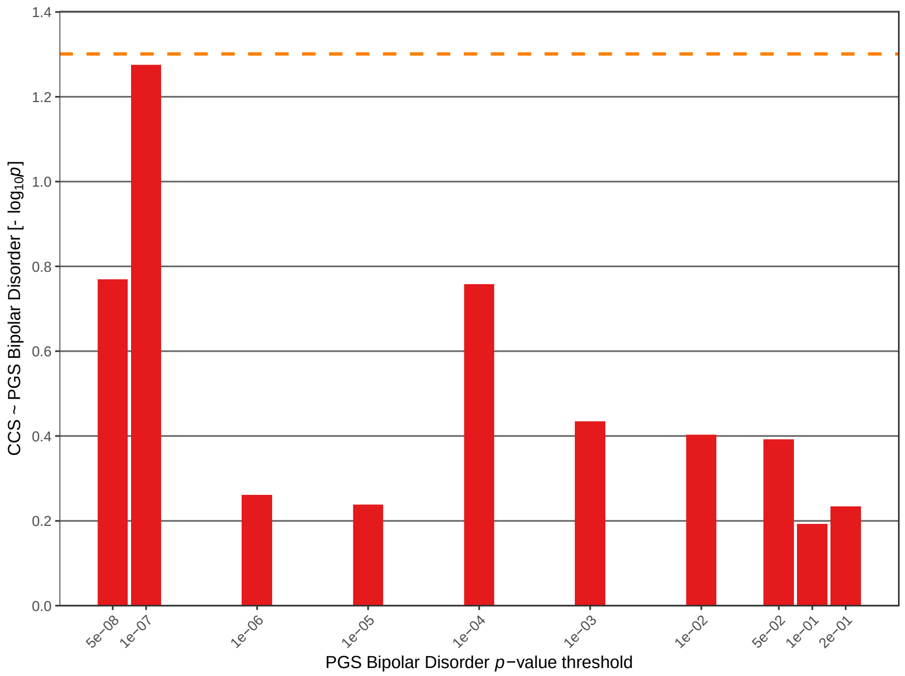


**D:** Analysis of PGS based on the MDD GWAS as training data.

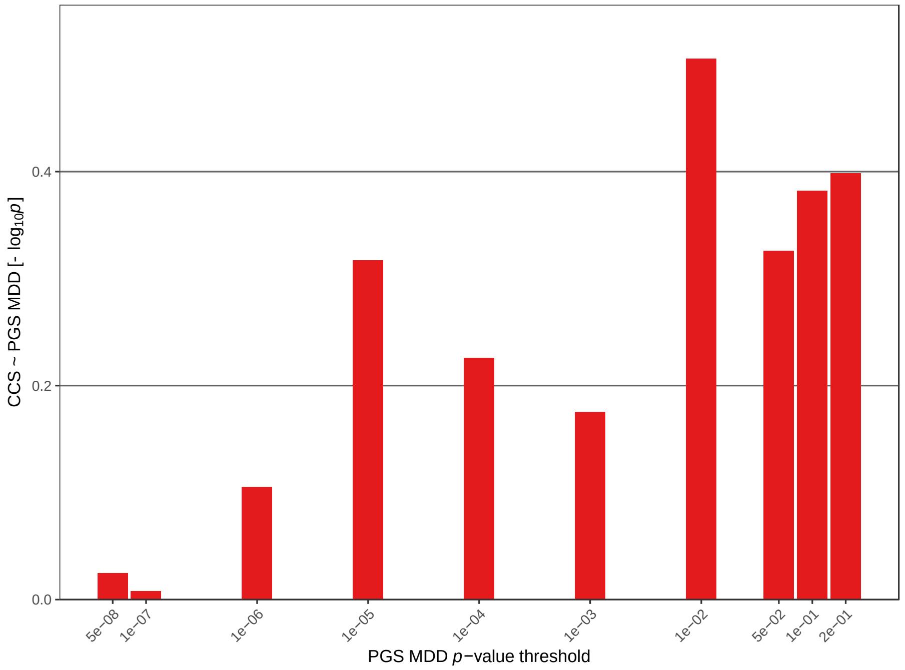

**E:** Analysis of PGS based on the Schizophrenia GWAS as training data.

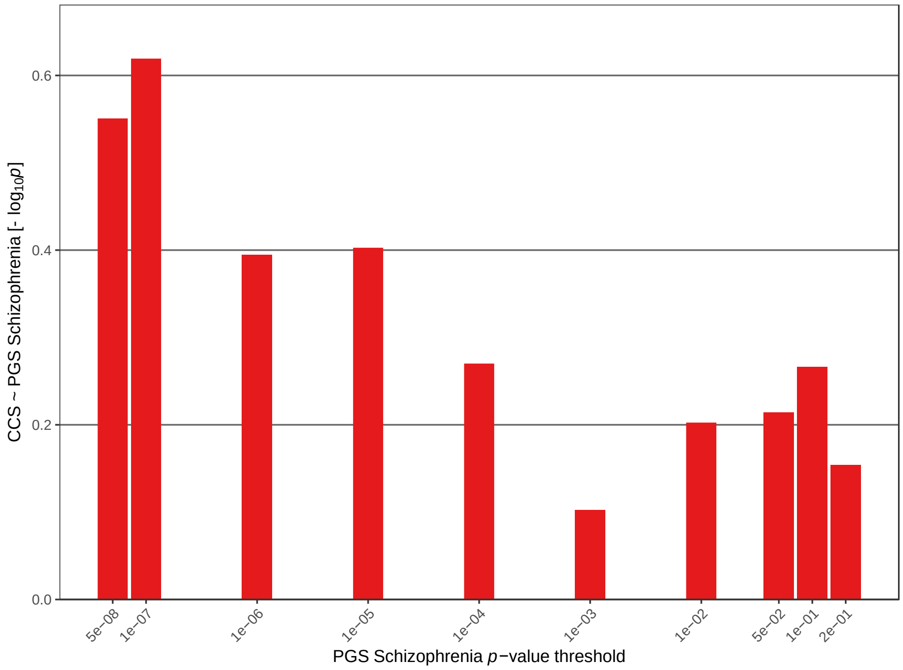


**F:** Analysis of PGS based on the bvFTD GWAS as training data.

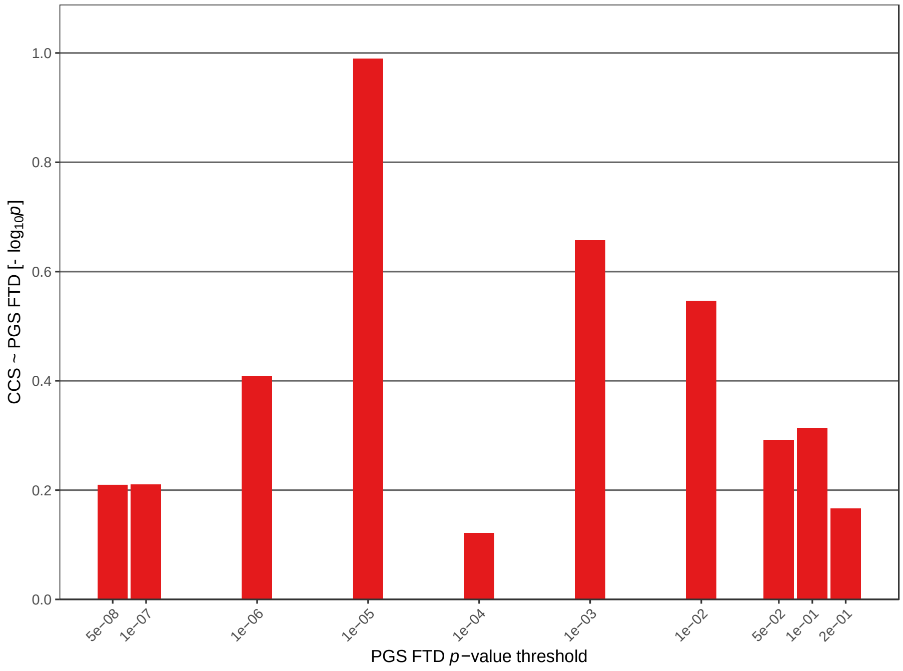

**G:** Analysis of PGS based on the epigenetic aging acceleration (EAA; all brain regions) GWAS as training data.

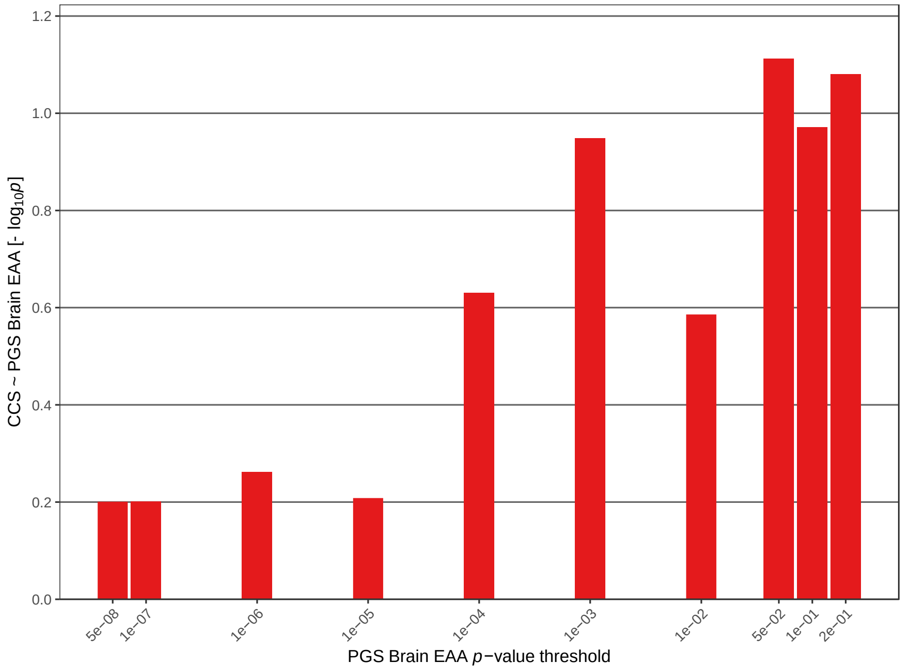


**H:** Analysis of PGS based on the EAA (prefrontal cortex (PFCTX)) GWAS as training data.

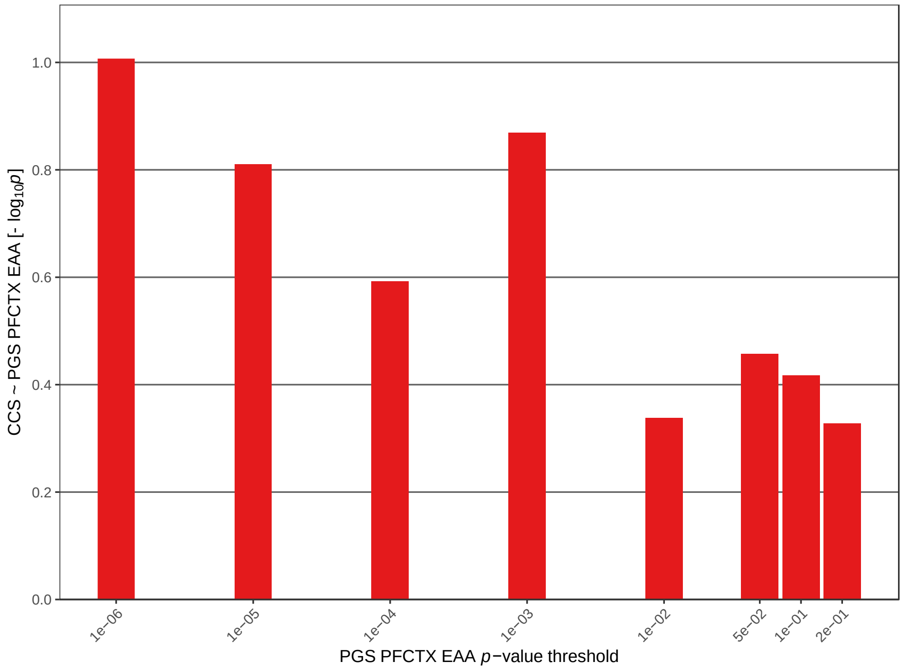

**I:** Analysis of PGS based on the neuronal proportion (prefrontal cortex) GWAS as training data.

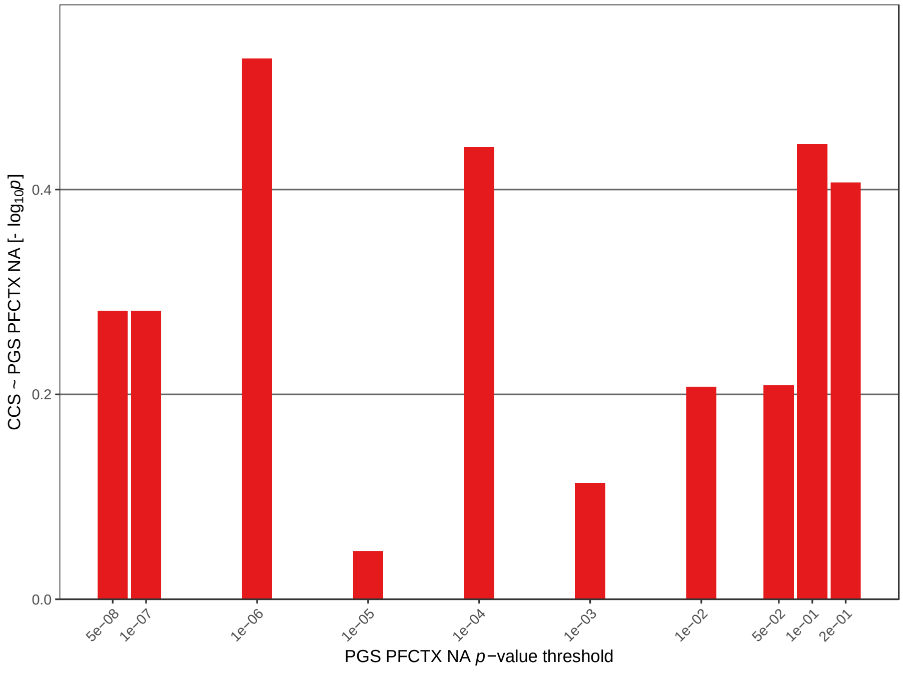


**J:** Analysis of PGS based on the longevity 85 GWAS as training data.

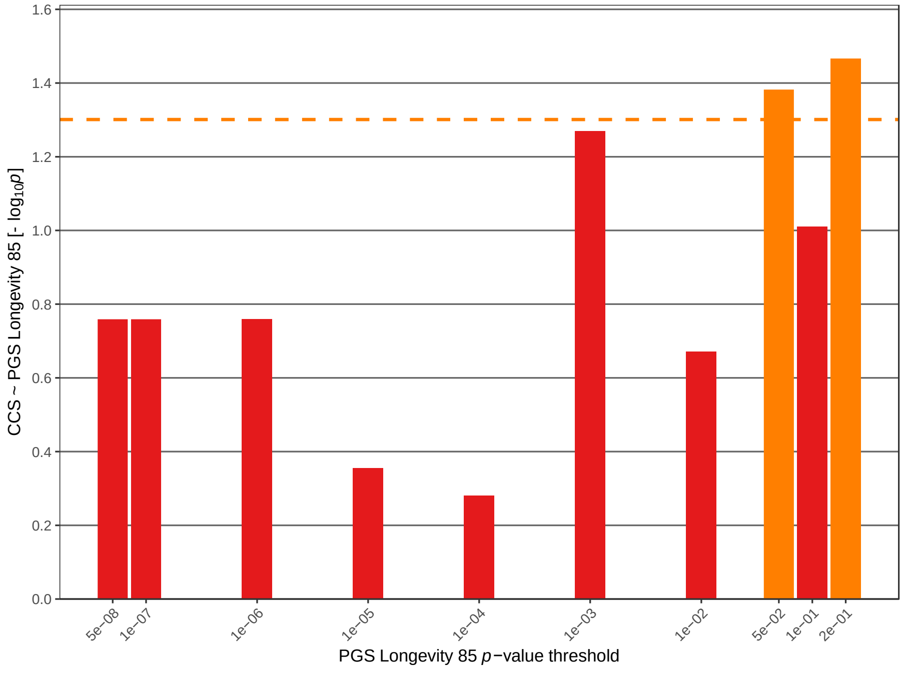

**K:** Analysis of PGS based on the longevity 90 GWAS as training data.

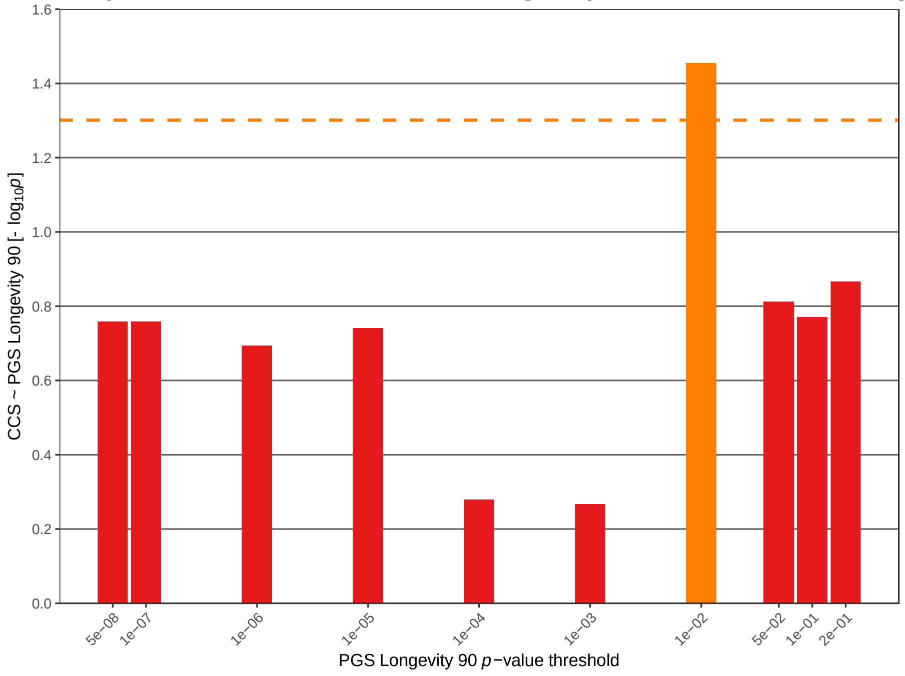

**Figure S7:** Box-whisker plot of the CCS per diagnosis group in the FOR2107 cohort sorted by the median CCS.
BD-II: Bipolar Disorder (BD) type 2, BD-I: BD type 1, BD NOS: not otherwise specified BD, SCZ: Schizophrenia, SZA: Schizoaffective Disorder.

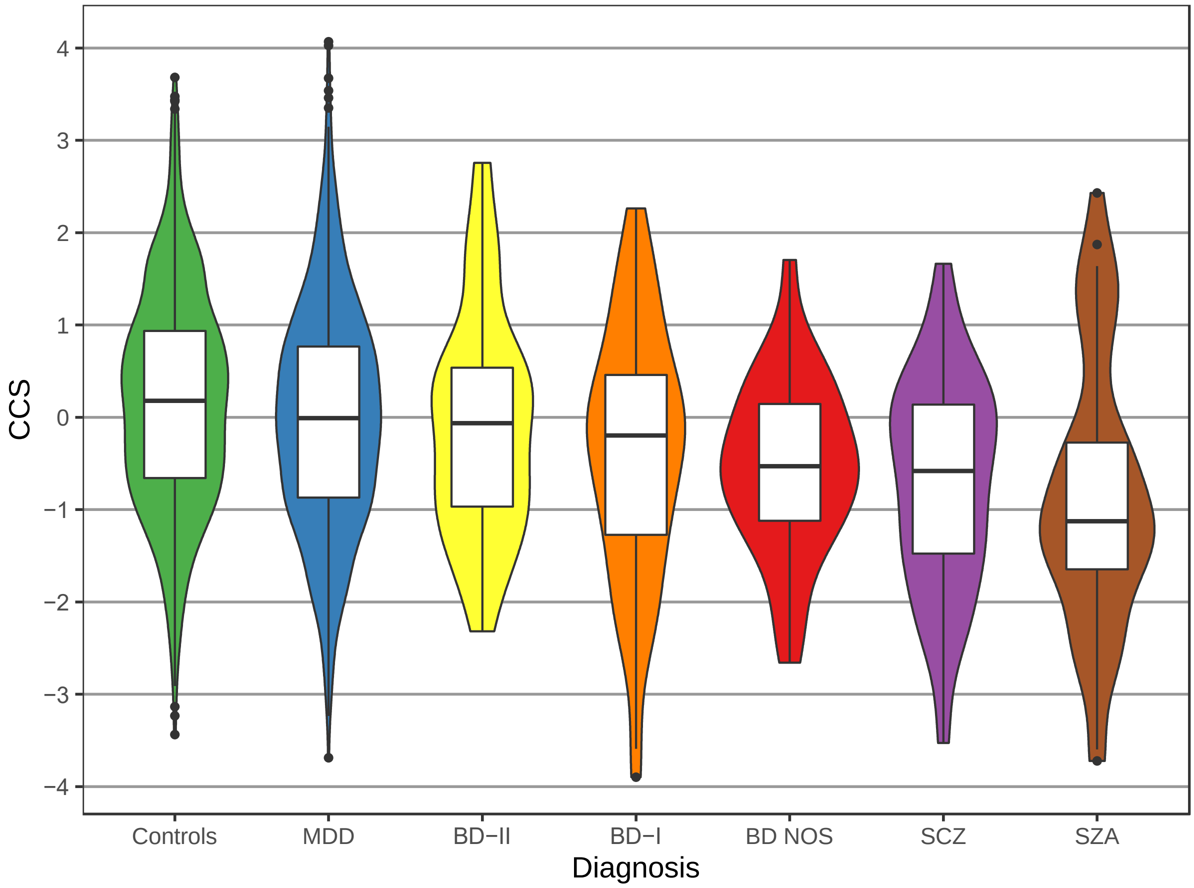

**Figure S8:** Rank-rank hypergeometric overlap maps.
Comparison of CCS-associated and bvFTD, EAA, and longevity GWAS-associated variants. For further details see Table S11.

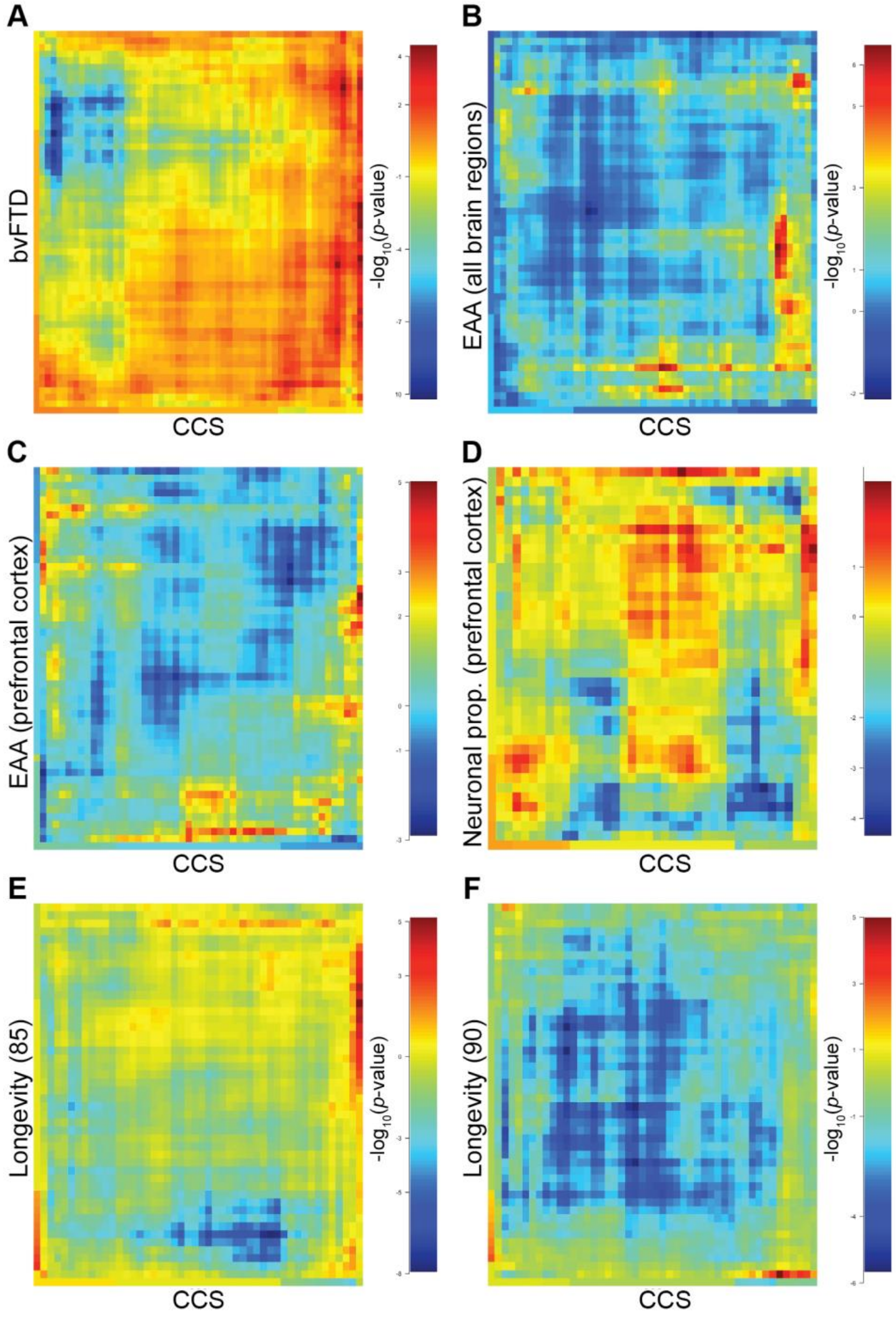

#### IFGC members and affiliations

Raffaele Ferrari (UCL, Department of Molecular Neuroscience, Russell Square House, 9-12 Russell Square House, London, WC1B 5EH), Dena G Hernandez (Laboratory of Neurogenetics, National Institute on Aging, National Institutes of Health, Building 35, Room 1A215, 35 Convent Drive, Bethesda, MD 20892, USA; Reta Lila Weston Research Laboratories, Department of Molecular Neuroscience, UCL Institute of Neurology, London WC1N 3BG, UK), Michael A Nalls (Laboratory of Neurogenetics, National Institute on Aging, National Institutes of Health Building 35, Room 1A215, 35 Convent Drive, Bethesda, MD 20892, USA), Jonathan D Rohrer (Reta Lila Weston Research Laboratories, Department of Molecular Neuroscience, UCL Institute of Neurology, Queen Square, London WC1N 3BG, UK; Dementia Research Centre, Department of Neurodegenerative Disease, UCL Institute of Neurology), Adaikalavan Ramasamy (Reta Lila Weston Research Laboratories, Department of Molecular Neuroscience, UCL Institute of Neurology, London WC1N 3BG, UK; Department of Medical and Molecular Genetics, King’s College London Tower Wing, Guy’s Hospital, London SE1 9RT, UK; The Jenner Institute, University of Oxford, Roosevelt Drive, Oxford OX3 7BQ, UK), John BJ Kwok (Neuroscience Research Australia, Sydney, NSW 2031, Australia; School of Medical Sciences, University of New South Wales, Sydney, NSW 2052, Australia), Carol Dobson-Stone (Neuroscience Research Australia, Sydney, NSW 2031, Australia; School of Medical Sciences, University of New South Wales, Sydney, NSW 2052, Australia), William S Brooks (Neuroscience Research Australia, Sydney, NSW 2031, Australia; Prince of Wales Clinical School, University of New South Wales, Sydney, NSW 2052, Australia), Peter R Schofield (Neuroscience Research Australia, Sydney, NSW 2031, Australia; School of Medical Sciences, University of New South Wales, Sydney, NSW 2052, Australia), Glenda M Halliday (Neuroscience Research Australia, Sydney, NSW 2031, Australia; School of Medical Sciences, University of New South Wales, Sydney, NSW 2052, Australia), John R Hodges (Neuroscience Research Australia, Sydney, NSW 2031, Australia; School of Medical Sciences, University of New South Wales, Sydney, NSW 2052, Australia), Olivier Piguet (Neuroscience Research Australia, Sydney, NSW 2031, Australia; School of Medical Sciences, University of New South Wales, Sydney, NSW 2052, Australia), Lauren Bartley (Neuroscience Research Australia, Sydney, NSW 2031, Australia), Elizabeth Thompson (South Australian Clinical Genetics Service, SA Pathology (at Women's and Children's Hospital), North Adelaide, SA 5006, Australia; Department of Paediatrics, University of Adelaide, Adelaide, SA 5000, Australia), Isabel Hernández (Research Center and Memory Clinic of Fundació ACE, Institut Català de Neurociències Aplicades, Barcelona, Spain), Agustín Ruiz (Research Center and Memory Clinic of Fundació ACE, Institut Català de Neurociències Aplicades, Barcelona, Spain), Mercè Boada (Research Center and Memory Clinic of Fundació ACE, Institut Català de Neurociències Aplicades, Barcelona, Spain), Barbara Borroni (Neurology Clinic, University of Brescia, Brescia, Italy), Alessandro Padovani (Neurology Clinic, University of Brescia, Brescia, Italy), Carlos Cruchaga (Department of Psychiatry, Washington University, St. Louis, MO, USA; Hope Center, Washington University School of Medicine, St. Louis, MO, USA), Nigel J Cairns (Hope Center, Washington University School of Medicine, St. Louis, MO, USA; Department of Pathology and Immunology, Washington University, St. Louis, MO, USA), Luisa Benussi (Molecular Markers Laboratory, IRCCS Istituto Centro San Giovanni di Dio Fatebenefratelli,Brescia, Italy), Giuliano Binetti (MAC Memory Clinic, IRCCS Istituto Centro San Giovanni di Dio Fatebenefratelli, Brescia, Italy), Roberta Ghidoni

(Molecular Markers Laboratory, IRCCS Istituto Centro San Giovanni di Dio Fatebenefratelli, Brescia, Italy), Gianluigi Forloni (Biology of Neurodegenerative Disorders, IRCCS Istituto di Ricerche Farmacologiche, "Mario Negri", Milano, Italy), Diego Albani (Biology of Neurodegenerative Disorders, IRCCS Istituto di Ricerche Farmacologiche "Mario Negri", Milano, Italy), Daniela Galimberti (University of Milan, Milan, Italy; Fondazione Cà Granda, IRCCS Ospedale Maggiore Policlinico, via F. Sforza 35, 20122, Milan, Italy), Chiara Fenoglio (University of Milan, Milan, Italy; Fondazione Cà Granda, IRCCS Ospedale Maggiore Policlinico, via F. Sforza 35, 20122, Milan, Italy), Maria Serpente (University of Milan, Milan, Italy; Fondazione Cà Granda, IRCCS Ospedale Maggiore Policlinico, via F. Sforza 35, 20122, Milan, Italy), Elio Scarpini (University of Milan, Milan, Italy; Fondazione Cà Granda, IRCCS Ospedale Maggiore Policlinico, via F. Sforza 35, 20122, Milan, Italy), Jordi Clarimón (Memory Unit, Neurology Department and Sant Pau Biomedical Research Institute, Hospital de la Santa Creu i Sant Pau, Universitat Autònoma de Barcelona, Barcelona, Spain; Center for Networker Biomedical Research in Neurodegenerative Diseases (CIBERNED), Madrid, Spain), Alberto Lleó (Memory Unit, Neurology Department and Sant Pau Biomedical Research Institute, Hospital de la Santa Creu i Sant Pau, Universitat Autònoma de Barcelona, Barcelona, Spain; Center for Networker Biomedical Research in Neurodegenerative Diseases (CIBERNED), Madrid, Spain), Rafael Blesa (Memory Unit, Neurology Department and Sant Pau Biomedical Research Institute, Hospital de la Santa Creu i Sant Pau, Universitat Autònoma de Barcelona, Barcelona, Spain; Center for Networker Biomedical Research in Neurodegenerative Diseases (CIBERNED), Madrid, Spain), Maria Landqvist Waldö (Unit of Geriatric Psychiatry, Department of Clinical Sciences, Lund University, Lund, Sweden), Karin Nilsson (Unit of Geriatric Psychiatry, Department of Clinical Sciences, Lund University, Lund, Sweden), Christer Nilsson (Clinical Memory Research Unit, Department of Clinical Sciences, Lund University, Lund, Sweden), Ian RA Mackenzie (Department of Pathology and Laboratory Medicine, University of British Columbia, Vancouver, Canada), Ging-Yuek R Hsiung (Division of Neurology, University of British Columbia, Vancouver, Canada), David MA Mann (Institute of Brain, Behaviour and Mental Health, University of Manchester, Salford Royal Hospital, Stott Lane, Salford, M6 8HD, UK), Jordan Grafman (Rehabilitation Institute of Chicago, Departments of Physical Medicine and Rehabilitation, Psychiatry, and Cognitive Neurology & Alzheimer's Disease Center; Feinberg School of Medicine, Northwestern University, Chicago, USA; Department of Psychology, Weinberg College of Arts and Sciences, Northwestern University, Chicago, USA), Christopher M Morris (Newcastle Brain Tissue Resource, Institute for Ageing, Newcastle University, NE4 5PL, Newcastle upon Tyne, UK; Newcastle University, Institute of Neuroscience and Institute for Ageing, Campus for Ageing and Vitality, NE4 5PL, Newcastle upon Tyne, UK; Institute of Neuroscience, Newcastle University Medical School, Framlington Place NE2 4HH, Newcastle upon Tyne, UK), Johannes Attems (Newcastle University, Institute of Neuroscience and Institute for Ageing, Campus for Ageing and Vitality, NE4 5PL, Newcastle upon Tyne, UK), Timothy D Griffiths (Institute of Neuroscience, Newcastle University Medical School, Framlington Place, NE2 4HH, Newcastle upon Tyne, UK), Ian G McKeith (Newcastle University, Institute of Neuroscience and Institute for Ageing, Campus for Ageing and Vitality, Newcastle University, NE4 5PL, Newcastle upon Tyne, UK), Alan J Thomas (Newcastle University, Institute of Neuroscience and Institute for Ageing, Campus for Ageing and Vitality, NE4 5PL, Newcastle upon Tyne, UK), Pietro Pietrini (IMT School for Advanced Studies, Lucca, Lucca, Italy), Edward D Huey (Taub Institute, Departments of Psychiatry and Neurology, Columbia University,

630 West 168th Street New York, NY 10032), Eric M Wassermann (Behavioral Neurology Unit, National Insititute of Neurological Disorders and Stroke, National Insititutes of Health, 10 CENTER DR MSC 1440, Bethesda, MD 20892-1440), Atik Baborie (Department of Laboratory Medicine & Pathology, Walter Mackenzie Health Sciences Centre, 8440 - 112 St, University of Alberta Edmonton, Alberta T6G 2B7, Canada), Evelyn Jaros (Newcastle University, Institute for Ageing and Health, Campus for Ageing and Vitality, NE4 5PL, Newcastle upon Tyne, UK), Michael C Tierney (Behavioral Neurology Unit, National Insititute of Neurological Disorders and Stroke, National Insititutes of Health, 10 CENTER DR MSC 1440, Bethesda, MD 20892-1440), Pau Pastor (Center for Networker Biomedical Research in Neurodegenerative Diseases (CIBERNED), Madrid, Spain; Neurogenetics Laboratory, Division of Neurosciences, Center for Applied Medical Research, Universidad de Navarra, Pamplona, Spain; Department of Neurology, Clínica Universidad de Navarra, University of Navarra School of Medicine, Pamplona, Spain), Cristina Razquin (Neurogenetics Laboratory, Division of Neurosciences, Center for Applied Medical Research, Universidad de Navarra, Pamplona, Spain), Sara Ortega-Cubero (Center for Networker Biomedical Research in Neurodegenerative Diseases (CIBERNED), Madrid, Spain; Neurogenetics Laboratory, Division of Neurosciences, Center for Applied Medical Research, Universidad de Navarra, Pamplona, Spain), Elena Alonso (Neurogenetics Laboratory, Division of Neurosciences, Center for Applied Medical Research, Universidad de Navarra, Pamplona, Spain), Robert Perneczky (Neuroepidemiology and Ageing Research Unit, School of Public Health, Faculty of Medicine, The Imperial College of Science, Technology and Medicine, London W6 8RP, UK; West London Cognitive Disorders Treatment and Research Unit, West London Mental Health Trust, London TW8 8 DS, UK; Department of Psychiatry and Psychotherapy, Technische Universität München, Munich, 81675 Germany), Janine Diehl-Schmid (Department of Psychiatry and Psychotherapy, Technische Universität München, Munich, 81675 Germany), Panagiotis Alexopoulos (Department of Psychiatry and Psychotherapy, Technische Universität München, Munich, 81675 Germany), Alexander Kurz (Department of Psychiatry and Psychotherapy, Technische Universität München, Munich, 81675 Germany), Innocenzo Rainero (Neurology I, Department of Neuroscience, University of Torino, Italy, A.O. Città della Salute e della Scienza di Torino, Torino, Italy), Elisa Rubino (Neurology I, Department of Neuroscience, University of Torino, Italy, A.O. Città della Salute e della Scienza di Torino, Torino, Italy), Lorenzo Pinessi (Neurology I, Department of Neuroscience, University of Torino, Italy, A.O. Città della Salute e della Scienza di Torino, Torino, Italy), Ekaterina Rogaeva (Tanz Centre for Research in Neurodegenerative Diseases, University of Toronto 60 Leonard Street, Toronto, Ontario, Canada, M5T 2S8), Peter St George-Hyslop (Tanz Centre for Research in Neurodegenerative Diseases, University of Toronto, 60 Leonard Street, Toronto, Ontario, Canada, M5T 2S8; Cambridge Institute for Medical Research, and the Department of Clinical Neurosciences, University of Cambridge, Hills Road, Cambridge, UK CB2 0XY), Giacomina Rossi (Division of Neurology V and Neuropathology, Fondazione IRCCS Istituto Neurologico Carlo Besta, 20133 Milano, Italy), Fabrizio Tagliavini (Division of Neurology V and Neuropathology, Fondazione IRCCS Istituto Neurologico Carlo Besta, 20133 Milano, Italy), Giorgio Giaccone (Division of Neurology V and Neuropathology, Fondazione IRCCS Istituto Neurologico Carlo Besta, 20133 Milano, Italy), James B Rowe (Cambridge University Department of Clinical Neurosciences, Cambridge, CB2 0SZ, UK; MRC Cognition and Brain Sciences Unit, Cambridge, CB2 7EF, UK; Behavioural and Clinical Neuroscience Institute, Cambridge, CB2 3EB, UK),

Johannes CM Schlachetzki (University of California San Diego, Department of Cellular & Molecular Medicine, 9500 Gilman Drive, La Jolla, CA, 92093), James Uphill (MRC Prion Unit, Department of Neurodegenerative Disease, UCL Institute of Neurology, Queen Square House, Queen Square, London, WC1N 3BG), John Collinge (MRC Prion Unit, Department of Neurodegenerative Disease, UCL Institute of Neurology Queen Square House, Queen Square, London, WC1N 3BG), Simon Mead (MRC Prion Unit, Department of Neurodegenerative Disease, UCL Institute of Neurology, Queen Square House, Queen Square, London, WC1N 3BG), Adrian Danek (Neurologische Klinik und Poliklinik, Ludwig-Maximilians-Universität, Munich, Germany, German Center for Neurodegenerative Diseases (DZNE)), Vivianna M Van Deerlin (University of Pennsylvania Perelman School of Medicine, Department of Pathology and Laboratory Medicine, Philadelphia, PA, USA), Murray Grossman (University of Pennsylvania Perelman School of Medicine, Department of Neurology and Penn Frontotemporal Degeneration Center, Philadelphia, PA, USA), John Q Trojanowski (University of Pennsylvania Perelman School of Medicine, Department of Pathology and Laboratory Medicine, Philadelphia, PA, USA), Julie van der Zee (Neurodegenerative Brain Diseases group, VIB-UAntwerp Center of Molecular Neurology, Antwerp, Belgium; Laboratory of Neurogenetics, Institute Born-Bunge, University of Antwerp, Antwerp, Belgium), Christine Van Broeckhoven (Neurodegenerative Brain Diseases group, VIB-UAntwerp Center of Molecular Neurology, Antwerp, Belgium; Laboratory of Neurogenetics, Institute Born-Bunge, University of Antwerp, Antwerp, Belgium), Stefano F Cappa (Neurorehabilitation Unit, Dept. Of Clinical Neuroscience, Vita-Salute University and San Raffaele Scientific Institute, Milan, Italy), Isabelle Leber (Inserm, UMR_S975, CRICM; UPMC Univ Paris 06, UMR_S975; CNRS UMR 7225, F-75013, Paris, France; AP-HP, Hôpital de la Salpêtrière, Département de neurologie-centre de références des démences rares, F-75013, Paris, France), Didier Hannequin (Service de Neurologie, Inserm U1079, CNR-MAJ, Rouen University Hospital, Rouen, France), Véronique Golfier (Service de neurologie, CH Saint Brieuc, France), Martine Vercelletto (Service de neurologie, CHU Nantes, France), Alexis Brice (Inserm, UMR_S975, CRICM; UPMC Univ Paris 06, UMR_S975; CNRS UMR 7225, F-75013, Paris, France; AP-HP, Hôpital de la Salpêtrière, Département de neurologie-centre de références des démences rares, F-75013, Paris, France), Benedetta Nacmias (Department of Neurosciences, Psychology, Drug Research and Child Health (NEUROFARBA) University of Florence, Florence, Italy), Sandro Sorbi (Department of Neurosciences, Psychology, Drug Research and Child Health (NEUROFARBA), University of Florence and IRCCS "Don Carlo Gnocchi" Firenze, Florence, Italy), Silvia Bagnoli (Department of Neurosciences, Psychology, Drug Research and Child Health (NEUROFARBA), University of Florence, Florence, Italy), Irene Piaceri (Department of Neurosciences, Psychology, Drug Research and Child Health (NEUROFARBA), University of Florence, Florence, Italy), Jørgen E Nielsen (Danish Dementia Research Centre, Neurogenetics Clinic, Department of Neurology, Rigshospitalet, Copenhagen University Hospital, Copenhagen, Denmark; Department of Cellular and Molecular Medicine, Section of Neurogenetics, The Panum Institute, University of Copenhagen, Copenhagen, Denmark), Lena E Hjermind (Danish Dementia Research Centre, Neurogenetics Clinic, Department of Neurology, Rigshospitalet, Copenhagen University Hospital, Copenhagen, Denmark; Department of Cellular and Molecular Medicine, Section of Neurogenetics, The Panum Institute, University of Copenhagen, Copenhagen, Denmark), Matthias Riemenschneider (Saarland University Hospital, Department for Psychiatry & Psychotherapy, Kirrberger Str.1, Bld.90, 66421 Homburg/Saar, Germany;

Saarland University, Laboratory for Neurogenetics, Kirrberger Str.1, Bld.90, 66421 Homburg/Saar, Germany), Manuel Mayhaus (Saarland University, Laboratory for Neurogenetics, Kirrberger Str.1, Bld.90, 66421 Homburg/Saar, Germany), Bernd Ibach (University Regensburg, Department of Psychiatry, Psychotherapy and Psychosomatics, Universitätsstr. 84, 93053 Regensburg, Germany), Gilles Gasparoni (Saarland University, Laboratory for Neurogenetics, Kirrberger Str.1, Bld.90, 66421 Homburg/Saar, Germany), Sabrina Pichler (Saarland University, Laboratory for Neurogenetics, Kirrberger Str.1, Bld.90, 66421 Homburg/Saar, Germany), Wei Gu (Saarland University, Laboratory for Neurogenetics, Kirrberger Str.1, Bld.90, 66421 Homburg/Saar, Germany; Luxembourg Centre For Systems Biomedicine (LCSB), University of Luxembourg 7, avenue des Hauts-Fourneaux, 4362 Esch- sur-Alzette, Luxembourg), Martin N Rossor (Dementia Research Centre, Department of Neurodegenerative Disease, UCL Institute of Neurology, Queen Square, London, WC1N 3BG), Nick C Fox (Dementia Research Centre, Department of Neurodegenerative Disease, UCL Institute of Neurology Queen Square, London, WC1N 3BG), Jason D Warren (Dementia Research Centre, Department of Neurodegenerative Disease, UCL Institute of Neurology, Queen Square, London, WC1N 3BG), Maria Grazia Spillantini (University of Cambridge, Department of Clinical Neurosciences, John Van Geest Brain Repair Centre, Forvie Site, Robinson way, Cambridge CB2 0PY), Huw R Morris (UCL, Department of Molecular Neuroscience, Russell Square House, 9-12 Russell Square House, London, WC1B 5EH), Patrizia Rizzu (German Center for Neurodegenerative Diseases-Tübingen, Otfried Muellerstrasse 23, Tuebingen 72076, Germany), Peter Heutink (German Center for Neurodegenerative Diseases-Tübingen, Otfried Muellerstrasse 23, Tuebingen 72076, Germany), Julie S Snowden (Institute of Brain, Behaviour and Mental Health, Faculty of Medical and Human Sciences, University of Manchester, Manchester, UK), Sara Rollinson (Institute of Brain, Behaviour and Mental Health, Faculty of Medical and Human Sciences, University of Manchester, Manchester, UK), Anna Richardson (Salford Royal Foundation Trust, Faculty of Medical and Human Sciences, University of Manchester, Manchester, UK), Alexander Gerhard (Institute of Brain, Behaviour and Mental Health, The University of Manchester, 27 Palatine Road, Withington, Manchester, M20 3LJ, UK), Amalia C Bruni (Regional Neurogenetic Centre, ASPCZ, Lamezia Terme, Italy), Raffaele Maletta (Regional Neurogenetic Centre, ASPCZ, Lamezia Terme, Italy), Francesca Frangipane (Regional Neurogenetic Centre, ASPCZ, Lamezia Terme, Italy), Chiara Cupidi (Regional Neurogenetic Centre, ASPCZ, Lamezia Terme, Italy), Livia Bernardi (Regional Neurogenetic Centre, ASPCZ, Lamezia Terme, Italy), Maria Anfossi (Regional Neurogenetic Centre, ASPCZ, Lamezia Terme, Italy), Maura Gallo (Regional Neurogenetic Centre, ASPCZ, Lamezia Terme, Italy), Maria Elena Conidi (Regional Neurogenetic Centre, ASPCZ, Lamezia Terme, Italy), Nicoletta Smirne (Regional Neurogenetic Centre, ASPCZ, Lamezia Terme, Italy), Rosa Rademakers (Department of Neuroscience, Mayo Clinic Jacksonville, 4500 San Pablo Road, Jacksonville, FL 32224), Matt Baker (Department of Neuroscience, Mayo Clinic Jacksonville, 4500 San Pablo Road, Jacksonville, FL 32224), Dennis W Dickson (Department of Neuroscience, Mayo Clinic Jacksonville, 4500 San Pablo Road, Jacksonville, FL 32224), Neill R Graff-Radford (Department of Neurology, Mayo Clinic Jacksonville, 4500 San Pablo Road, Jacksonville, FL 32224), Ronald C Petersen (Department of Neurology, Mayo Clinic Rochester, 2001st street SW Rochester MN 5905), David Knopman (Department of Neurology, Mayo Clinic Rochester, 2001st street SW Rochester MN 5905), Keith A Josephs (Department of Neurology, Mayo Clinic Rochester, 2001st street SW Rochester MN 5905), Bradley F

Boeve (Department of Neurology, Mayo Clinic Rochester, 2001st street SW Rochester MN 5905), Joseph E Parisi (Department of Pathology, Mayo Clinic Rochester, 2001st street SW Rochester MN 5905), William W Seeley (Department of Neurology, Box 1207, University of California, San Francisco, CA 94143, USA), Bruce L Miller (Memory and Aging Center, Department of Neurology, University of California, San Francisco, CA 94158, USA), Anna M Karydas (Memory and Aging Center, Department of Neurology, University of California, San Francisco, CA 94158, USA), Howard Rosen (Memory and Aging Center, Department of Neurology, University of California, San Francisco, CA 94158, USA), John C van Swieten (Department of Neurology, Erasmus Medical Centre, Rotterdam, The Netherlands; Department of Medical Genetics, VU university Medical Centre, Amsterdam, The Netherlands), Elise GP Dopper (Department of Neurology, Erasmus Medical Centre, Rotterdam, The Netherlands), Harro Seelaar (Department of Neurology, Erasmus Medical Centre, Rotterdam, The Netherlands), Yolande AL Pijnenburg (Alzheimer Centre and department of neurology, VU University medical centre, Amsterdam, The Netherlands), Philip Scheltens (Alzheimer Centre and department of neurology, VU University medical centre, Amsterdam, The Netherlands), Giancarlo Logroscino (Department of Basic Medical Sciences, Neurosciences and Sense Organs of the "Aldo Moro" University of Bari, Bari, Italy), Rosa Capozzo (Department of Basic Medical Sciences, Neurosciences and Sense Organs of the "Aldo Moro" University of Bari, Bari, Italy), Valeria Novelli (Medical Genetics Unit, Fondazione Policlinico Universitario A. Gemelli, Rome, Italy), Annibale A Puca (Cardiovascular Research Unit, IRCCS Multimedica, Milan, Italy; Department of Medicine and Surgery, University of Salerno, Baronissi (SA), Italy), Massimo Franceschi (Neurology Dept, IRCCS Multimedica, Milan, Italy), Alfredo Postiglione (Department of Clinical Medicine and Surgery, University of Naples Federico II, Naples, Italy), Graziella Milan (Geriatric Center Frullone- ASL Napoli 1 Centro, Naples, Italy), Paolo Sorrentino (Geriatric Center Frullone- ASL Napoli 1 Centro, Naples, Italy), Mark Kristiansen (UCL Genomics, Institute of Child Health (ICH), UCL, London, UK), Huei-Hsin Chiang (Karolinska Institutet, Dept NVS, Alzheimer Research Center, Novum, SE-141 57, Stockholm, Sweden; Dept of Geriatric Medicine, Genetics Unit, M51, Karolinska University Hospital, SE-14186, Stockholm), Caroline Graff (Karolinska Institutet, Dept NVS, Alzheimer Research Center, Novum, SE-141 57, Stockholm, Sweden; Dept of Geriatric Medicine, Genetics Unit, M51, Karolinska University Hospital, SE-14186, Stockholm), Florence Pasquier (Univ Lille, Inserm 1171, DISTALZ, CHU 59000 Lille, France), Adeline Rollin (Univ Lille, Inserm 1171, DISTALZ, CHU 59000 Lille, France), Vincent Deramecourt (Univ Lille, Inserm 1171, DISTALZ, CHU 59000 Lille, France), Thibaud Lebouvier (Univ Lille, Inserm 1171, DISTALZ, CHU 59000 Lille, France), Dimitrios Kapogiannis (National Institute on Aging (NIA/NIH), 3001 S. Hanover St, NM 531, Baltimore, MD, 21230), Luigi Ferrucci (Clinical Research Branch, National Institute on Aging, Baltimore, MD, USA), Stuart Pickering-Brown (Institute of Brain, Behaviour and Mental Health, Faculty of Medical and Human Sciences, University of Manchester, Manchester, UK), Andrew B Singleton (Laboratory of Neurogenetics, National Institute on Aging, National Institutes of Health, Building 35, Room 1A215, 35 Convent Drive, Bethesda, MD 20892, USA), John Hardy (UCL, Department of Molecular Neuroscience, Russell Square House, 9-12 Russell Square House, London, WC1B 5EH), Parastoo Momeni (Laboratory of Neurogenetics, Department of Internal Medicine, Texas Tech University Health Science Center, 4th street, Lubbock, Texas 79430, USA).

#### IFGC acknowledgments

Intramural funding from the National Institute of Neurological Disorders and Stroke (NINDS) and National Institute on Aging (NIA), the Wellcome/MRC Centre on Parkinson's disease, Alzheimer's Research UK (ARUK, Grant ARUK-PG2012-18) and by the office of the Dean of the School of Medicine, Department of Internal Medicine, at Texas Tech University Health Sciences Center. We thank Mike Hubank and Kerra Pearce at the Genomic core facility at the Institute of Child Health (ICH), University College of London (UCL), for assisting RF in performing Illumina genotyping experiments (FTD-GWAS genotyping). This study utilized the high-performance computational capabilities of the Biowulf Linux cluster at the National Institutes of Health, Bethesda, Md. (http://biowulf.nih.gov). North American Brain Expression Consortium (NABEC) - The work performed by the North American Brain Expression Consortium (NABEC) was supported in part by the Intramural Research Program of the National Institute on Aging, National Institutes of Health, part of the US Department of Health and Human Services; project number ZIA AG000932-04. In addition this work was supported by a Research Grant from the Department of Defense, W81XWH-09-2-0128. UK Brain Expression Consortium (UKBEC) - This work performed by the UK Brain Expression Consortium (UKBEC) was supported by the MRC through the MRC Sudden Death Brain Bank (C.S.), by a Project Grant (G0901254 to J.H. and M.W.) and by a Fellowship award (G0802462 to M.R.). D.T. was supported by the King Faisal Specialist Hospital and Research Centre, Saudi Arabia. Computing facilities used at King's College London were supported by the National Institute for Health Research (NIHR) Biomedical Research Centre based at Guy's and St Thomas' NHS Foundation Trust and King's College London. We would like to thank AROS Applied Biotechnology AS company laboratories and Affymetrix for their valuable input. RF's work is supported by Alzheimer's Society (grant number 284), UK; JBJK was supported by the National Health and Medical Resarch Council (NHMRC) Australia, Project Grants 510217 and 1005769; CDS was supported by NHMRC Project Grants 630428 and 1005769; PGS was supported by NHMRC Project Grants 510217 and 1005769 and acknowledges that DNA samples were prepared by Genetic Repositories Australia, supported by NHMRC Enabling Grant 401184; GMH was supported by NHMRC Research Fellowship 630434, Project Grant 1029538, Program Grant 1037746; JRH was supported by the Australian Research Council Federation Fellowship, NHMRC Project Grant 1029538, NHMRC Program Grant 1037746; OP was supported by NHMRC Career Development Fellowship 1022684, Project Grant 1003139. IH, AR and MB acknowledge the patients and controls who participated in this project and the Trinitat Port-Carbó and her family who are supporting Fundació ACE research programs. CC was supported by Grant P30- NS069329-01 and acknowledges that the recruitment and clinical characterization of research participants at Washington University were supported by NIH P50 AG05681, P01 AG03991, and P01 AG026276. LB and GB were supported by the Ricerca Corrente, Italian Ministry of Health; RG was supported by Fondazione CARIPLO 2009-2633, Ricerca Corrente, Italian Ministry of Health; GF was supported by Fondazione CARIPLO 2009-2633. ES was supported by the Italian Ministry of Health; CF was supported by Fondazione Cariplo; MS was supported from the Italian Ministry of Health (Ricerca Corrente); MLW was supported by Government funding of clinical research within NHS Sweden (ALF); KN was supported by Thure Carlsson Foundation; CN was supported by Swedish Alzheimer Fund. IRAM and GYRH were supported by CIHR (grant 74580) PARF (grant C06-01). JG was supported by the NINDS intramural research funds for FTD research. CMM was supported by Medical Research Council UK, Brains for Dementia Research, Alzheimer's Society, Alzheimer's Research UK, National Institutes for Health Research, Department of Health, Yvonne Mairy Bequest and acknowledges that tissue made available for this study was provided by the Newcastle Brain Tissue Resource, which was funded in part by grants G0400074 and G1100540 from the UK MRC, the Alzheimer's Research Trust and Alzheimer's Society through the Brains for Dementia Research Initiative and an NIHR Biomedical Research Centre Grant in Ageing and Health, and NIHR Biomedical Research Unit in Lewy Body Disorders. CMM was supported by the UK Department of Health and Medical Research Council and the Research was supported by the National Institute for Health Research Newcastle Biomedical Research Centre based at Newcastle Hospitals Foundation Trust and Newcastle University and acknowledges that the views expressed are those of the authors and not necessarily those of the NHS, the NIHR or the Department of Health; JA was supported by MRC, Dunhill Medical Trust, Alzheimer's Research UK; TDG was supported by Wellcome Trust Senior Clinical Fellow; IGM was supported by NIHR Biomedical Research Centre and Unit on Ageing Grants and acknowledges the National Institute for Health Research Newcastle Biomedical Research Centre based at Newcastle Hospitals Foundation Trust and Newcastle University. The views expressed are those of the author(s) and not necessarily those of the NHS, the NIHR or the Department of Health; AJT was supported by Medical Research Council, Alzheimer's Society, Alzheimer's Research UK, National Institutes for Health Research. EJ was supported by NIHR, Newcastle Biomedical Research Centre. PP, CR, SOC and EA were supported partially by FIMA (Foundation for Applied Medical Research); PP acknowledges Manuel Seijo-Martínez (Department of Neurology, Hospital do Salnés, Pontevedra, Spain), Ramon Rene, Jordi Gascon and Jaume Campdelacreu (Department of Neurology, Hospital de Bellvitge, Barcelona, Spain) for providing FTD DNA samples. RP, JDS, PA and AK were supported by German Federal Ministry of Education and Research (BMBF; grant number FKZ 01GI1007A – German FTLD consortium). IR was supported by Ministero dell'Istruzione, dell'Università e della Ricerca (MIUR) of Italy. PStGH was supported by the Canadian Institutes of Health Research, Wellcome Trust, Ontario Research Fund. FT was supported by the Italian Ministry of Health (ricerca corrente) and MIUR grant RBAP11FRE9; GR and GG were supported by the Italian Ministry of Health (ricerca corrente). JBR was supported by Camrbidge NIHR Biomedical Research Centre and Wellcome Trust (088324). JU, JC, SM were supported by the MRC Prion Unit core funding and acknowledge MRC UK, UCLH Biomedical Research Centre, Queen Square Dementia BRU; SM acknowledges the work of John Beck, Tracy Campbell, Gary Adamson, Ron Druyeh, Jessica Lowe, Mark Poulter. AD acknowledges the work of Benedikt Bader and of Manuela Neumann, Sigrun Roeber, Thomas Arzberger and Hans Kretzschmar†; VMVD and JQT were supported by Grants AG032953, AG017586 and AG010124; MG was supported by Grants AG032953, AG017586, AG010124 and NS044266; VMVD acknowledges EunRan Suh, PhD for assistance with sample handling and Elisabeth McCarty-Wood for help in selection of cases; JQT acknowledges Terry Schuck, John Robinson and Kevin Raible for assistance with neuropathological evaluation of cases. CVB and the Antwerp site were in part funded by the MetLife Foundation for Medical Research Award (to CVB); the Belgian Science Policy Office (BELSPO) Interuniversity Attraction Poles program; the Flemish Government initiated Methusalem Excellence Program (to CVB); the Flemish government initiated Impulse Program on Networks for Dementia Research (VIND); the Research Foundation Flanders (FWO) and the University of Antwerp Research Fund. CVB and JvdZ acknowledge the neurologists S

Engelborghs, PP De Deyn, A Sieben, R Vandenberghe and the neuropathologist JJ Martin for the clinical and pathological diagnoses. CVB and JvdZ further thank the personnel of the Neuromics Support Facility of the VIB Center for Molecular Neurology and the Antwerp Biobank of the Institute Born-Bunge for their expert support. IL and AB were supported by the program “Investissements d’avenir” ANR-10-IAIHU-06 and acknowledges the contribution of The French research network on FTLD/FTLD-ALS for the contribution in samples collection. BN is founded by Fondazione Cassa di Risparmio di Pistoia e Pescia (grant 2014.0365), SS is founded by the Cassa di Risparmio di Firenze (grant 2014.0310) and a grant from Ministry of Health n° RF-2010-2319722. JEN was supported by the Novo Nordisk Foundation, Denmark. MR was supported by the German National Genome Network (NGFN); German Ministry for Education and Research Grant Number 01GS0465. JDR, MNR, NCF and JDW were supported by an MRC programme grant and the Dementia Platform UK, the NIHR Queen Square Dementia Biomedical Research Unit (BRU) and the Leonard Wolfson Experimental Neurology Centre. MGS was supported by MRC grant n G0301152, Cambridge Biomedical Research Centre and acknowledges Mrs K Westmore for extracting DNA. HM was supported by the Motor Neuron Disease Association (Grant 6057). RR was supported by P50 AG016574, R01 NS080882, R01 NS065782, P50 NS72187 and the Consortium for Frontotemporal Dementia; DWD was supported by P50NS072187, P50AG016574, State of Florida Alzheimer Disease Initiative, & CurePSP, Inc.; NRGR, JEP, RCP, DK, BFB were supported by P50 AG016574; KAJ was supported by R01 AG037491; WWS was supported by NIH AG023501, AG019724, Consortium for Frontotemporal Dementia Research; BLM was supported by P50AG023501, P01AG019724, Consortium for FTD Research; HR was supported by AG032306. JCvS was supported by Stichting Dioraphte Foundation (11 02 03 00), Nuts Ohra Foundation (0801-69), Hersenstichting Nederland (BG 2010-02) and Alzheimer Nederland. CG and HHC acknowledge families, patients, clinicians including Dr Inger Nennesmo and Dr Vesna Jelic, Professor Laura Fratiglioni for control samples and Jenny Björkström, Håkan Thonberg, Charlotte Forsell, Anna-Karin Lindström and Lena Lilius for sample handling. CG was supported by Swedish Brain Power (SBP), the Strategic Research Programme in Neuroscience at Karolinska Institutet (StratNeuro), the regional agreement on medical training and clinical research (ALF) between Stockholm County Council and Karolinska Institutet, Swedish Alzheimer Foundation, Swedish Research Council, Karolinska Institutet PhD-student funding, King Gustaf V and Queen Victoria’s Free Mason Foundation. FP, AR, VD and FL acknowledge Labex DISTALZ. RF acknowledges the help and support of Mrs. June Howard at the Texas Tech University Health Sciences Center Office of Sponsored Programs for tremendous help in managing Material Transfer Agreement at TTUHSC.

#### FOR2107 acknowledgments

This work is part of the German multicenter consortium "Neurobiology of Affective Disorders. A translational perspective on brain structure and function", funded by the German Research Foundation (Deutsche Forschungsgemeinschaft DFG; Forschungsgruppe/Research Unit FOR2107).

Principal investigators (PIs) with respective areas of responsibility in the FOR2107 consortium are:
Work Package WP1, FOR2107/MACS cohort and brainimaging: Tilo Kircher (speaker FOR2107; DFG grant numbers KI 588/14-1, KI 588/14-2), Udo Dannlowski (co-speaker FOR2107; DA 1151/5-1, DA 1151/5-2), Axel Krug (KR 3822/5-1, KR 3822/7-2), Igor Nenadic (NE 2254/1-2), Carsten Konrad (KO 4291/3-1). WP2, animal phenotyping: Markus Wöhr (WO 1732/4-1, WO 1732/4-2), Rainer Schwarting (SCHW 559/14-1, SCHW 559/14-2). WP3, miRNA: Gerhard Schratt (SCHR 1136/3-1, 1136/3-2). WP4, immunology, mitochondriae: Judith Alferink (AL 1145/5-2), Carsten Culmsee (CU 43/9-1, CU 43/9-2), Holger Garn (GA 545/5-1, GA 545/7-2). WP5, genetics: Marcella Rietschel (RI 908/11-1, RI 908/11-2), Markus Nöthen (NO 246/10-1, NO 246/10-2), Stephanie Witt (WI 3439/3-1, WI 3439/3-2). WP6, multimethod data analytics: Andreas Jansen (JA 1890/7-1, JA 1890/7-2), Tim Hahn (HA 7070/2-2), Bertram Müller-Myhsok (MU1315/8-2), Astrid Dempfle (DE 1614/3-1, DE 1614/3-2). CP1, biobank: Petra Pfefferle (PF 784/1-1, PF 784/1-2), Harald Renz (RE 737/20-1, 737/20-2). CP2, administration. Tilo Kircher (KI 588/15-1, KI 588/17-1), Udo Dannlowski (DA 1151/6-1), Carsten Konrad (KO 4291/4-1).

Data access and responsibility: All PIs take responsibility for the integrity of the respective study data and their components. All authors and coauthors had full access to all study data.
Acknowledgements and members by Work Package (WP):
WP1: Henrike Bröhl, Katharina Brosch, Bruno Dietsche, Rozbeh Elahi, Jennifer Engelen, Sabine Fischer, Jessica Heinen, Svenja Klingel, Felicitas Meier, Tina Meller, Julia-Katharina Pfarr, Kai Ringwald, Torsten Sauder, Simon Schmitt, Frederike Stein, Annette Tittmar, Dilara Yüksel (Dept. of Psychiatry, Marburg University). Mechthild Wallnig, Rita Werner (Core-Facility Brainimaging, Marburg University). Carmen Schade-Brittinger, Maik Hahmann (Coordinating Centre for Clinical Trials, Marburg). Michael Putzke (Psychiatric Hospital, Friedberg). Rolf Speier, Lutz Lenhard (Psychiatric Hospital, Haina). Birgit Köhnlein (Psychiatric Practice, Marburg). Peter Wulf, Jürgen Kleebach, Achim Becker (Psychiatric Hospital Hephata, Schwalmstadt-Treysa). Ruth Bär (Care facility Bischoff, Neukirchen). Matthias Müller, Michael Franz, Siegfried Scharmann, Anja Haag, Kristina Spenner, Ulrich Ohlenschläger (Psychiatric Hospital Vitos, Marburg). Matthias Müller, Michael Franz, Bernd Kundermann (Psychiatric Hospital Vitos, Gießen). Christian Bürger, Katharina Dohm, Fanni Dzvonyar, Verena Enneking, Stella Fingas, Katharina Förster, Janik Goltermann, Dominik Grotegerd, Hannah Lemke, Susanne Meinert, Nils Opel, Ronny Redlich, Jonathan Repple, Kordula Vorspohl, Bettina Walden, Dario Zaremba (Dept. of Psychiatry, University of Münster). Harald Kugel, Jochen Bauer, Walter Heindel, Birgit Vahrenkamp (Dept. of Clinical Radiology, University of Münster). Gereon Heuft, Gudrun Schneider (Dept. of Psychosomatics and Psychotherapy, University of Münster). Thomas Reker (LWL-Hospital Münster). Gisela Bartling (IPP Münster). Ulrike Buhlmann (Dept. of Clinical Psychology, University of Münster).

WP5: Helene Dukal, Christine Hohmeyer, Lennard Stütz, Viola Lahr, Fabian Streit, Josef Frank, Lea Sirignano (Dept. of Genetic Epidemiology, Central Institute of Mental Health, Medical Faculty Mannheim, Heidelberg University). Stefanie Heilmann-Heimbach, Stefan Herms, Per Hoffmann (Institute of Human Genetics, University of Bonn, School of Medicine & University Hospital Bonn). Andreas J. Forstner (Institute of Human Genetics, University of Bonn, School of Medicine & University Hospital Bonn; Centre for Human Genetics, Marburg University).
WP6: Anastasia Benedyk, Miriam Bopp, Roman Keßler, Maximilian Lückel, Verena Schuster, Christoph Vogelbacher (Dept. of Psychiatry, Marburg University). Jens Sommer, Olaf Steinsträter (Core-Facility Brainimaging, Marburg University). Thomas W.D. Möbius (Institute of Medical Informatics and Statistics, Kiel University).
CP1: Julian Glandorf, Fabian Kormann, Arif Alkan, Fatana Wedi, Lea Henning, Alena Renker, Karina Schneider, Elisabeth Folwarczny, Dana Stenzel, Kai Wenk, Felix Picard, Alexandra Fischer, Sandra Blumenau, Beate Kleb, Doris Finholdt, Elisabeth Kinder, Tamara Wüst, Elvira Przypadlo, Corinna Brehm (Comprehensive Biomaterial Bank Marburg, Marburg University).
The FOR2107 cohort project (WP1) was approved by the Ethics Committees of the Medical Faculties, University of Marburg (AZ: 07/14) and University of Münster (AZ: 2014-422-b-S).
The study was supported by the German Federal Ministry of Education and Research (BMBF), through the Integrated Network IntegraMent, under the auspices of the e:Med programme (grants 01ZX1314A/01ZX1614A to MMN; 01ZX1314G/01ZX1614G to MR; 01ZX1614J to BMM) through grants 01EE1406C to MR and 01EE1409C to MR and SHW, and through ERA-NET NEURON, “SynSchiz - Linking synaptic dysfunction to disease mechanisms in schizophrenia - a multilevel investigation“ (01EW1810 to MR) and BMBF grants 01EE1409C and 01EE1406C to MR and SHW.

#### MPIP acknowledgments

We wish to acknowledge Rosa Schirmer, Elke Schreiter, Reinhold Borschke and Ines Eidner for image acquisition and data preparation, Anna Olynyik for data quality control, and Benno Pütz, Nazanin Karbalai, Darina Czamara, and Bertram Müller-Myhsok for distributed computing support. We thank Dorothee P. Auer and Florian Holsboer for initiation of the RUD study. The MARS study was supported by a grant of the Exzellenz-Stiftung of the Max Planck Society. This work has also been funded by the BMBF in the framework of the National Genome Research Network (NGFN) (FKZ 01GS0481). The funders had no role in study design, data collection and analysis, decision to publish, or preparation of the manuscript.
